# Individual tree-based vs pixel-based approaches to mapping forest functional traits and diversity by remote sensing

**DOI:** 10.1101/2022.07.10.499231

**Authors:** Zhaoju Zheng, Yuan Zeng, Meredith C. Schuman, Hailan Jiang, Bernhard Schmid, Michael E. Schaepman, Felix Morsdorf

**Affiliations:** Remote Sensing Laboratories, Department of Geography, University of Zurich, Zurich CH-8057, Switzerland; State Key Laboratory of Remote Sensing Science, Aerospace Information Research Institute, Chinese Academy of Sciences, Beijing 100101, China; University of Chinese Academy of Sciences, Beijing 100049, China; State Key Laboratory of Remote Sensing Science, Faculty of Geographical Science, Beijing Normal University, Beijing 100875, China

**Keywords:** LiDAR, Imaging spectroscopy, Functional traits, Individual-tree approach, Pixel-based approach, Forest functional diversity

## Abstract

Trait-based approaches, focusing on the functional characteristics of vascular plants in a community, have been increasingly used in plant ecology and biodiversity research. Compared with traditional field survey (which typically samples individual trees), remote sensing enables quantifying functional traits over large contiguous areas, but assigning trait values to biological units such as species and individuals is difficult with pixel-based approaches. We used a subtropical forest landscape in China to compare an approach based on LiDAR-delineated individual tree crowns (ITCs) with a pixel-based approach for assessing functional traits from remote sensing data. We compared trait distributions, trait–trait relationships and functional diversity metrics obtained by the two approaches at changing grain and extent. We found that morphological traits derived from airborne laser scanning showed more differences between ITC- and pixel-based approaches than physiological traits estimated by imaging spectroscopy data. Pixel sizes approximating average tree crowns yielded similar results as ITCs, but 95th quantile height and foliage height diversity tended to be overestimated and leaf area index underestimated relative to ITC-based values. With increasing pixel size, the differences to ITC-based trait values became larger and less trait variance was captured, indicating information loss. The consistency of ITC- and pixel-based functional richness measures also decreased with increasing pixel grain, and changed with the observed extent for functional diversity monitoring. We conclude that whereas ITC-based approaches in principle allow partitioning of variation between individuals, genotypes and species, at high resolution, pixel-based approaches come close to this and can be suitable for assessing ecosystem-scale trait variation by weighting individuals and species according to coverage.

## 1. Introduction

Plant functional traits, often defined as the morphological, physiological or phenological features of vascular plants associated with individual fitness and ecological strategies, are sensitive indicators of ecological processes and ecosystem functioning (Homolová et al., 2013; Violle et al., 2007). With the emergence of functional biogeography (Violle et al., 2014), functional traits and trait-based functional diversity have received increasing attention in biodiversity and global-change research. Functional diversity can capture both inter- and intra- specific trait variability and be characterized by single or multiple traits using variance- or distance-based metrics (Laliberté and Legendre, 2010; Petchey and Gaston, 2002). How well functional diversity could explain ecosystem functioning depends on the traits and niches filled by organisms present in the ecosystem (Cadotte et al., 2011; Díaz and Cabido, 2001). As the most important global repositories of terrestrial biodiversity, forests provide vital ecosystem functions and services (Liang et al., 2016; Reich, 2012). Therefore, quantifying functional traits efficiently and accurately is essential for monitoring the diversity and functioning of forest communities.

In recent decades, the number of observational datasets (Kattge et al., 2020; Weigelt et al., 2020) and global maps (Butler et al., 2017; Moreno-Martínez et al., 2018) of plant traits has been constantly growing. However, the consistency and representativeness of the local samplings and the resolution of the global maps are still limited (Jetz et al., 2016). Therefore, filling data gaps with detailed information in trait variation is critical for ecological and biodiversity research. Complementary to field measurements, remote sensing provides an efficient way to obtain spatially contiguous trait data over large areas (Cawse-Nicholson et al., 2021; Gamon et al., 2019; Skidmore et al., 2021; Torabzadeh et al., 2014). For example, airborne light detection and ranging (LiDAR, also known as laser scanning) can penetrate through gaps between leaves and capture the vertical, horizontal and inner structure of the canopy accurately (Coops et al., 2016; Valbuena et al., 2020). Forest morphological traits such as height, fractional cover, leaf area index (LAI) and crown diameter can be directly derived from LiDAR data at either plot or tree level (Duncanson et al., 2014; Guo et al., 2017; Morsdorf et al., 2006, 2004; Næsset, 2002; Popescu and Wynne, 2004). Besides, imaging spectroscopy provides continuous spectral information relating to light interactions with chemical and physical properties of leaves or canopies (Gamon et al., 2019; Schaepman et al., 2015; Ustin and Gamon, 2010). It has been widely used to retrieve foliar physiological traits, such as leaf pigments (Blackburn, 2007; Feilhauer et al., 2015; Ustin et al., 2009), leaf mass per area (Asner et al., 2011; Singh et al., 2015), foliar nitrogen, carbon, phosphorus (Asner et al., 2015; Serbin et al., 2014; Wang et al., 2016b), leaf water (Casas et al., 2014), lignin and cellulose contents (Kokaly et al., 2009) at leaf and canopy levels (Serbin and Townsend, 2020).

An important aspect of remote sensing-based approaches is that they generally assess community-weighted means of functional traits (Ma et al., 2020) without identifying the biological units such as species, genotypes and individuals that express these traits. Trait variation within plant communities is caused by genetic and environmental variation among these biological units as well as environmental variation within individuals (Baur and Schmid, 1996). Thus, a potential disadvantage of remotely-sensed datasets compared with the ground-based datasets could be the lack of species- and individual-level information about functional traits. To assess the degree of this potential disadvantage, it would be necessary to obtain remote sensing data for individual tree crowns (ITCs), and then use these for a comparison with the commonly used remotely sensed pixel-level data.

The combination of airborne LiDAR and imaging spectroscopy has allowed to mapping morphological and physiological traits of forests at different spatial units. A typical spatial unit is a quadratic grid cell or pixel, which leads to continuous trait maps at high (e.g. 1 m, 2 m, 6 m) or medium (e.g. 30 m) spatial resolutions (Asner et al., 2015; Coops et al., 2016; Schneider et al., 2017; Wang et al., 2020). A pixel, as the basic spatial unit of a digital image, is a geometric concept based on imaging and projection on a detector array, while field-based surveys collect trait data of individual species and trees. To obtain the functional trait of a plot, community-weighted means of functional traits are calculated, e.g. using species abundance as a weighting variable (Abelleira Martínez et al., 2016). The plot (or the tree community) is therefore the unit corresponding to the concept of pixels. For images with coarse spatial resolution, spectrally mixed pixels (due to the presence of multiple species and canopy gaps) can diminish the variability between pixels (Hacker et al., 2022). However, as plots or pixels get smaller, it can be assumed that the difference between individuals and community becomes smaller. Although a pixel typically does not assign information to individuals, small pixels are more likely to include only one species than do large pixels. New object-based image-analysis methods can aggregate pixels of high-resolution images to larger objects such as forest patches, eventually making it possible to work with individual tree crowns instead of pixels (Dechesne et al., 2017; Wu and Zhang et al., 2020). Furthermore, advances in LiDAR sensor technology and segmentation algorithms (Ferraz et al., 2016; Kaartinen et al., 2012; Wang et al., 2016a) increasingly enable the identification of ITCs for which information such as species identity (Shi et al., 2018; Torabzadeh et al., 2019), functional traits (Ferreira et al., 2018; Marconi et al., 2021; Zheng et al., 2021) or even genotype identity (Guillén-Escribà et al., 2021) may then be extracted.

Whereas the ITC-based approach can capture fine-scale individual-level trait variation, repeated acquisition of very-high-resolution airborne data over large areas will often make it too costly. On the other hand, there is a high demand for assessing the state and change of Earth’s plant diversity globally (Jetz et al., 2016), where the relatively coarse pixel of satellite remote sensing are generally used as a basic observation unit. Since most satellite sensors’ pixel sizes are much larger than tree crowns, retrieving traits for large pixels based on in-situ plot-level traits would average individual or species trait variation. However, it remains largely unknown how much fine-scale trait variation could be captured with increasing pixel size and at what level functional diversity can be observed from coarser pixels.

Scale is a fundamental concept in ecology (Levin, 1992; Wu et al., 2006) and is an essential practical consideration in both remote sensing and traditional biodiversity studies (Anderson, 2018; Gamon et al., 2020; Steinbauer et al., 2012). Given the various concepts of the term “scale”, we confine our discussion of scale to the spatial domain, focusing on two important components: grain (the unit of measurement, e.g. individual trees or pixels at different spatial resolutions) and extent (the geographical area measured) (Wiens, 1989). Previous studies in grassland have shown that changing the grain of observations could affect perceived diversity patterns and may alter associated conclusions in ecological processes of interest (Gholizadeh et al., 2019, 2018; Wang et al., 2018). As grain increases, a larger proportion of trait variation is averaged within the grain (grid cell or pixel), and between-grain heterogeneity decreases (Hay et al., 2001; Wiens, 1989). Besides, increasing spatial extent typically incorporates greater environmental heterogeneity as well as a larger range of trait values in the community, which has also been demonstrated by positive functional richness–area relationships in many studies (Durán et al., 2019; Karadimou et al., 2016; Schneider et al., 2017; Zheng et al., 2021).

In this study, we combine airborne LiDAR and imaging spectroscopy data to estimate both ITC- and pixel-based functional traits and functional diversity in a species-rich subtropical forest. We compare the distribution of ITC- and pixel-based functional traits at changing grain and extent and further investigate how much variability of functional traits would be lost when pixel size increases. Finally, we explore how well ITC-based functional diversity correlates with pixel-based functional diversity and discuss the suitability of both approaches.

## 2. Materials

### 2.1. Study area

This study was conducted in a subtropical evergreen and deciduous broad-leaved mixed forest in Xingshan county, Hubei province, China. A square area of 3 × 3 km (center coordinates: 31° 19′ N, 110° 29′ E) situated on the southern slope of Mt. Shennongjia was selected as the study area, with an altitude ranging from 1124 m to 1922 m a.s.l. (Fig. 1). Due to the complex species composition (approximately 20 dominant tree species), topography and microclimate variation in this mountain area, the forest canopy is highly heterogeneous, both horizontally and vertically. The forested areas are dominated by evergreen broad-leaved species (*Cyclobalanopsis multinervis*, *Cyclobalanopsis oxyodon* and *Lithocarpus glaber*), deciduous broad-leaved species (*Betula luminifera*, *Fagus engleriana*, *Platycarya strobilacea*, *Quercus aliena*, *Quercus serrata* and *Sorbus folgneri*) and coniferous species (*Cunninghamia lanceolata*, *Larix kaempferi* and *Pinus massoniana*) (Zeng et al., 2008; Zheng et al., 2021).

**Fig. 1.**
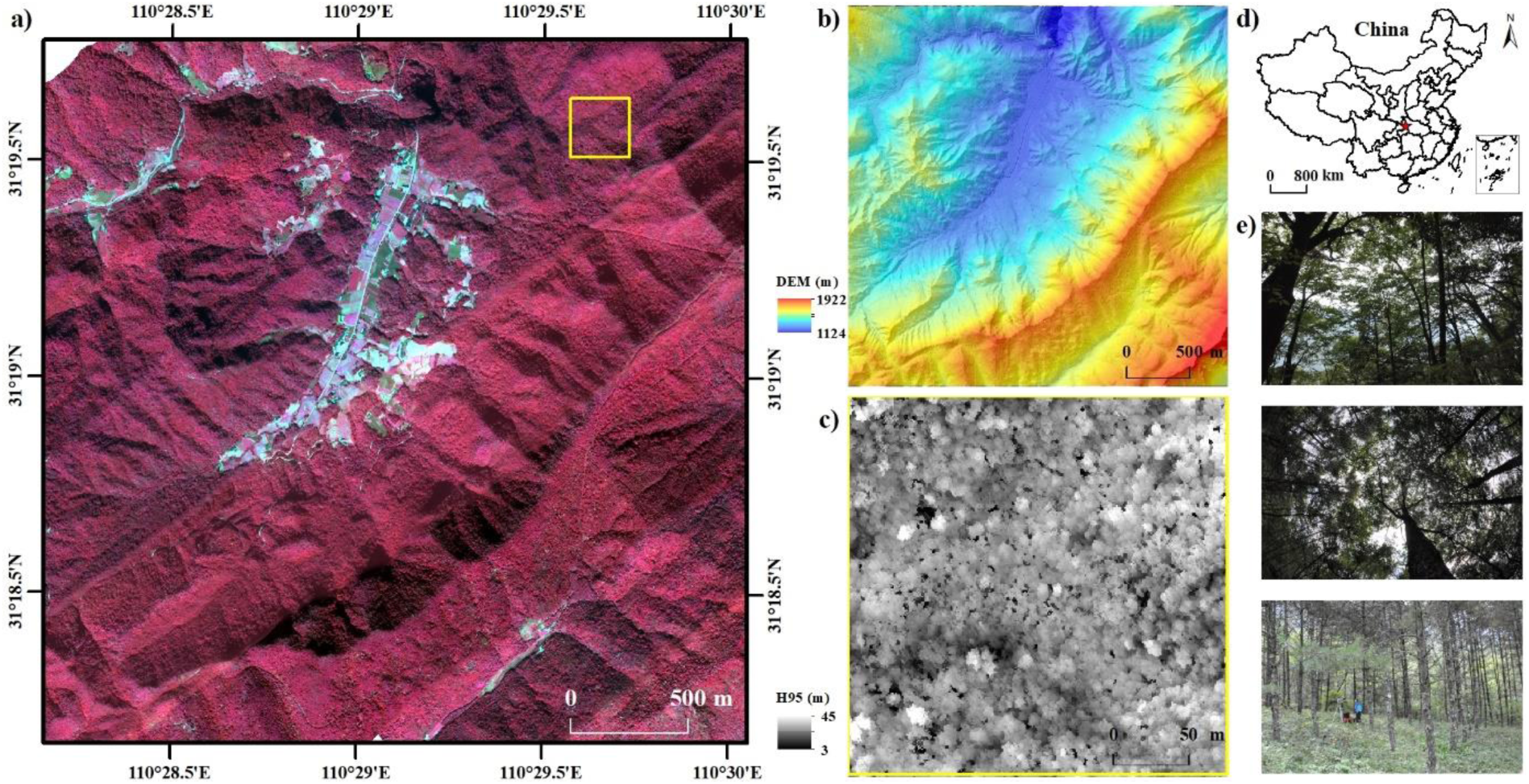
Overview of the study area and remote sensing data. (a) Study area (3 × 3 km) imaged with imaging spectroscopy data (Red: 810 nm, Green: 660 nm, Blue: 550 nm) at 1 m ground pixel size; the yellow square marks the 250 × 250 m subregion from which the histograms in Fig. 6 and Supp Fig. S6 were computed. (b) LiDAR-derived digital elevation model (DEM) of the study area and (c) 95th quantile height (H_95_) in the subregion at 1 m ground pixel size. (d) The studied subtropical forest is located in Hubei province of central China (the red star) and (e) shows in-situ photos of main forest types.

### 2.2. Airborne remote sensing data and preprocessing

The small-footprint LiDAR data were acquired during leaf-on conditions (October 2013) using a Leica ALS70-HP system mounted on a Yun-5 turboprop aircraft. The average flying altitude of 1800 m above ground level along with 0.15 mrad laser beam divergence led to a footprint diameter of approximately 0.27 m. The overlapping flight strips and the maximum scan angle of ±17° from the nadir led to a mean point density of 9.05 points m^−2^ (5.73 pulses m^−2^) in the study area. Multiple echo recordings (1–5 returns) were supplied as LAS 1.3 format files, with a basic noise filtering and classification of ground and non-ground returns already applied using Terrasolid software (Terrasolid, Helsinki, Finland) by the vendor (SHHANGYAO Inc., Shanghai, China). The DEM was constructed based on the classified ground points and further used for point clouds normalization. The elevation-normalized LiDAR point clouds were used to derive morphological traits for ITCs and pixels as described below (Section 3.2). ITCs were segmented from a 0.2-m rasterized canopy height model (CHM) using a morphological crown control-based watershed algorithm (Zhao et al., 2014). This ITC segmentation approach was used and validated in our previous study (Zheng et al., 2021) and the processing steps were also summarized here in Supp Text S1.

The airborne imaging spectroscopy campaign was carried out on 11 October and 13 October 2013 from 10:30 to 13:30 local time under clear sky conditions. Two sensors were mounted on the Pushbroom Hyperspectral Imager-3 (PHI-3) instrument, recording hyperspectral data in visible and near-infrared (0.44–1.0 µm) to shortwave-infrared (1.0–2.5 µm) spectral ranges with spectral resolutions of 3 nm and 5 nm, respectively. The instrument was mounted on the Yun-5 aircraft that flew at 1500 m above ground level and the field of view was 40 degrees, resulting in a ground resolution of 1 m. The data were radiometrically calibrated to achieve at-sensor radiances and then geometrically co-registered to ensure appropriate geometric co-alignment of imaging spectroscopy and LiDAR data. The preprocessed PHI-3 radiance data were atmospherically and topographically corrected using the rugged terrain modules of ATCOR-4. The bidirectional reflectance distribution function (BRDF) correction model of ATCOR-4 was used to reduce illumination effects (Schläpfer et al., 2015) . Subsequently, the reflectance images were orthorectified. Finally, the spectral regions dominated by atmospheric water vapor (1326–1518 nm and 1823–2015 nm) were removed and the resulting spectra were resampled to 10 nm bandwidth using the spectral resampling tool from ENVI 5.5. These reflectance images were used to estimate physiological traits as described below (Section 3.2).

### 2.3. Ground-based measurements

Field data were collected on the ground in the flight area during September to October 2013 (46 plots) and September 2016 (16 plots). The coordinates of plot corners for each 30 × 30 m plot were determined by a differentially-corrected Global Position System (Trimble GeoXH3000 handheld GPS) with sub-meter precision. Each tree in the plots with a diameter at breast height (DBH) greater than 5 cm was recorded with the species name, tree height, crown base height and approximate crown diameters. The tree positions were accurately located (< 10 cm) in four of these plots by integrating the Real-Time Kinematic (RTK) GPS/GLONASS System with a Leica TC302 total station for ITC detection assessment. We first used a Trimble integrated GNSS system with HBCORS service to obtain geodetic coordinates of two control points with centimeter-level precision (≈ 2 cm) in an open area close to the plot by RTK fixed solutions. These two control points were used for the total station instrument stationing and orientation. Afterward, we surveyed trees in the plot to determine the coordinates of the base of tree stems based on the polar coordinate method (da Silva et al., 2018). A comparison of LiDAR-detected ITC locations and field-measured tree locations and the ITC detection accuracy (match rate, omission, commission) at these four plots are provided in Zheng et al. (2021). Here, we further assessed the ITC detection rate for different tree height groups in this study (Supp Fig. S1). In addition, we collected sunlit top-of-canopy leaves from 31 overstory trees with known GPS locations during the 2016 field campaign and measured the leaf carotenoid content, specific leaf area (SLA) and nitrogen concentration (Zhao et al., 2016) to validate the remotely-sensed ITC-based physiological traits (Zheng et al., 2021).

## 3. Methods

### 3.1. Overview of methodology

The ITC- and pixel-based forest functional trait diversity assessments had three main parts (Fig. 2). First, morphological and physiological traits selected based on our previous study (Zheng et al., 2021) were derived from the airborne LiDAR and imaging spectroscopy data using these two approaches (Section 3.2). For the ITC-based approach, functional traits were extracted and aggregated within corresponding LiDAR-detected ITCs. For the pixel-based approach, the LiDAR point clouds were aggregated on a predefined grid and the imaging spectroscopy data were resampled to the corresponding pixel size (see Supp Fig. S2 for schematic diagram). Then, the differences of ITC- and pixel-based traits in distribution and the trait variability explained by different pixel sizes were compared (Section 3.3). Finally, a multi-dimensional functional diversity metric was calculated, and how well ITC- and pixel-based assessments of forest functional diversity agreed was investigated (Section 3.4).

**Fig. 2.**
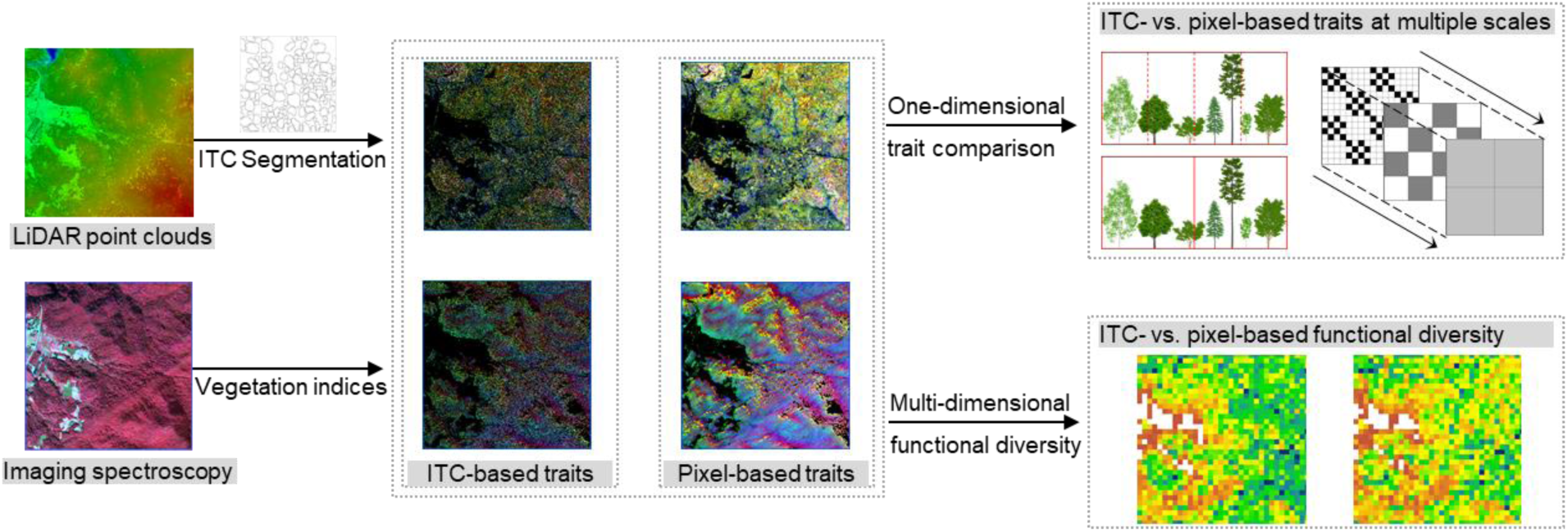
Flowchart of the processing steps.

### 3.2. ITC- and pixel-based functional trait estimations

An overview of the functional traits with corresponding biological function descriptions is provided in Table 1. For morphological traits, we estimated three commonly used height-, density- and layering-related structural parameters: 95th quantile height (H_95_), leaf area index (LAI, here more precisely plant area index because of the inclusion of non-leaf plant material) and foliage height diversity (FHD) from the elevation-normalized LiDAR point clouds within each ITC or pixel. We also set a normalized height threshold of 3 m to mask the sub-canopy trees or saplings and non-forest areas. H_95_ represents the height of a tree or forest canopy and can be calculated as the 95th quantile height of LiDAR first returns (Simonson et al., 2012). The LAI is defined as the projected leaf and branch area per unit ground area and can be retrieved from the gap fraction model based on the Beer-Lambert law (Richardson et al., 2009). FHD describes the canopy layering complexity and can be calculated following the formula of Shannon-Wiener index FHD = -∑p*_i_* ln p*_i_*, where p*_i_* is the proportion of horizontal vegetation in the *i*th layer (Clawges et al., 2008). Higher FHD values are expected with increasing canopy height extent and foliage arrangement evenness (Ehbrecht et al., 2016; MacArthur and MacArthur, 1961).

**Table 1.** Structural parameters and spectral indices used as indicators for morphological and physiological traits, respectively.

| Functional traits | Indicators | Function descriptions | References |
| --- | --- | --- | --- |
| Vertical height | 95th quantile height (H<sub>95</sub>) | affecting light resource allocation, forest productivity and successional dynamics | Simonson et al., 2012 |
| Horizontal density | Leaf area index (LAI) | leaf surface available for photosynthesis and transpiration and energy exchange | Richardson et al., 2009 |
| Structural complexity | Foliage height diversity (FHD) | characterizing foliage arrangement; related to light interception and tree growth | MacArthur and MacArthur, 1961 |
| Leaf carotenoids | Carotenoid reflectance index (CRI) | $CRI = 1/R_{510} - 1/R_{550}$, indicating the relative content of carotenoids per unit leaf area; important in photosynthesis and stress defense | Gitelson et al., 2002 |
| Specific leaf area | Simple ratio index (SR) | $SR = R_{800}/R_{680}$, indicating the ratio of leaf area to dry mass; reflecting carbon-gain strategies and leaf life span | Jordan, 1969 |
| Leaf nitrogen | Normalized difference nitrogen index (NDNI) | indicating the relative concentration of nitrogen per unit leaf mass; important for photosynthesis and metabolism | Serrano et al., 2002 |

For physiological traits, we calculated three spectral indices (VIs), namely carotenoid reflectance index (CRI), simple ratio index (SR) and normalized difference nitrogen index (NDNI) to represent leaf carotenoids, SLA and leaf nitrogen, which are ecologically-relevant functional traits related to forest growth, drought stress and productivity (Croft and Chen, 2018; Díaz et al., 2016; Wright et al., 2004). Since shaded and sunlit sides of tree crowns have different spectral features and shaded pixels generally suffer from a low signal-to-noise ratio (Clark et al., 2005, Trier et al., 2018), only fully illuminated canopy pixels were considered for VIs calculation by combining the shadow mask (CRI > 11.92) and non-vegetation mask (NDVI < 0.2). Additionally, we excluded non-forested areas (e.g. cropland, meadow) by applying a 3-m height threshold based on 3 m resolution H_95_. These masks were applied before ITC aggregation or pixel resampling since we wanted to focus on canopy information related to physiological trait variation for both approaches, and it would have been difficult to compare results if masks were applied afterward. For ITC-based traits, LiDAR-detected ITC polygons were used to extract the 1-m pixels belonging to each tree crown and their overlapping areas with the ITC polygon. The area-weighted mean VIs of these pixels were calculated to obtain ITC-based physiological trait values in IDL 8.6 (L3Harris Geospatial, Broomfield, CO, USA). For pixel-based traits, the 1-m VIs were resampled to predefined pixel sizes by pixel average (nanmean, python) to indicate physiological traits at different resolutions. To reduce the edge effect, we excluded ITCs and synthetic pixels that had more than half non-forested or shaded areas as determined during masking.

### 3.3. Comparison of ITC- and pixel-based traits

We calculated linear correlations to investigate how well ITC-based traits matched pixel-based traits at a resolution similar to the size of tree crowns (3 m side length). Since individual crown polygons did not necessarily fall within one pixel, we matched each ITC with the pixel closest to the treetop location. We further compared the frequency histograms (using the same bin width and anchor locations) of ITC- and pixel-based traits to investigate their distribution differences. Specifically, we used the median and the interquartile range (IQR) values to describe the center and spread of the trait distribution. We also assessed trait-to-trait relationships by calculating Pearson correlation coefficients (ρ) with two-tailed p-values to test whether trait-trait correlations generally hold between ITC- and pixel-based estimates.

In addition, we used the 250 × 250 m subregion (square in Fig. 1) to illustrate the distribution differences of functional traits derived from ITCs and pixels at different resolutions (3 m, 5 m, 10 m and 20 m). Besides the frequency histograms of ITC- and pixel-based traits, two-sample Kolmogorov-Smirnov tests (K-S test) and Mann-Whitney U tests were carried out to determine if the distributions of ITC- and pixel-based traits in the specific forest community were significantly different.

### 3.4. Scaling of functional traits and functional diversity

To assess how grain size and extent affect the variability of functional traits and the detected functional diversity patterns, we employed a scaling analysis by comparing the variation in functional traits from two aspects. In the first approach, we focused on single traits and grain changes. We calculated the coefficients of variation (CVs) of each functional trait over the study area and compared the between-unit variation in measured traits at increasing pixel sizes (3– 300 m). The between-unit trait variability can be defined as one-dimensional functional diversity or the extent of trait dissimilarity in a community because of the discrepancy between units (de Bello et al., 2009).

In the second approach, we assessed the multi-dimensional trait variation by functional richness (Villéger et al., 2008) . As a commonly-used functional diversity metric, functional richness refers to the functional niche occupied by the organisms in a community. It can be calculated as the convex hull volume of the multi-dimensional trait space occupied by ITCs or pixels at any given neighborhood extent (Schneider et al., 2017; Zheng et al., 2021). Since functional richness is relatively sensitive to outliers, we linearly scaled the traits from 0 to 1 using a min-max normalization after trimming the lower and upper (2nd to 98th percentile) extreme values (Schleuter et al., 2010). Then we calculated and mapped the morphological richness (Morph.FRic) and physiological richness (Phys.FRic) separately based on traits of ITCs and 3-m pixels in every 30 × 30 m and 100 × 100 m grid cell. We also calculated the pixel-wise Pearson correlation coefficients to investigate consistency between ITC- and 3-m pixel-based functional richness indices.

We also changed grain and extent together to investigate how well ITC- and pixel-based functional richness correlated at a given neighborhood extent when changing pixel size to the ones of high- to medium-resolution satellite imagery (3–30 m). To ensure a consistent number of analysis samples for all scales, we calculated the functional richness metrics at multiple spatial extents of concentric circles surrounding the same sample points. First, we generated 1000 random points within 300 m of the border and removed the points in non-forested areas (less than 10000 trees within a 250 m radius) as sample points. Then we calculated their Morph.FRic and Phys.FRic indices based on traits of ITCs or pixels within increasing neighborhood extents ranging from 9 to 294 m radii with a step of 7.5 m. The pixel sizes were not only fixed as 3 m, but also changed from 5 to 30 m with a step of 5 m. The minimum radius for functional richness calculations was set as at least 9 pixels involved in the circle. Pearson correlation coefficients were calculated to evaluate the consistency of ITC- and pixel-based functional richness indices at changing grain and extent.

Finally, we investigated the relationships between functional richness and area to examine the scale dependency of functional diversity and the spatial pattern of community assembly. Besides the observed ITC- and 3-m pixel-based functional richness–area curves for the sample points, we also calculated functional richness indices for these sample points based on the null model scenarios. The null models were developed by randomly reshuffling (randperm, Matlab) the locations of all ITCs and pixels within the study area. The comparison between the null model and observed data can also corroborate the robustness of the observed functional richness–area relationship and indicate if there is trait convergence or divergence in community assembly. We assessed the observed and null functional richness–area relationships for both ITCs and 3-m pixels and investigated whether these two approaches lead to the same community assembly patterns. All statistical analyses were carried out with SciPy library (Virtanen et al., 2020) in python 3.6.

## 4. Results

### 4.1. ITC- and pixel-based functional traits

The maps of morphological traits (Fig. 3) and physiological traits (Fig. 4) displayed different spatial patterns (See Supp Fig. S3 and Fig. S4 for maps of each trait). The trait validation performed in our previous study (Zheng et al., 2021) indicated that the LiDAR-derived morphological traits (R^2^ = 0.81 for LAI, R^2^ ≥ 0.90 for tree height; both P < 0.001) and spectral indices of physiological traits (R^2^ = 0.60 for carotenoids, R^2^ = 0.51 for SLA and R^2^ = 0.34 for nitrogen; all P < 0.001) correlated well with the field-measured individual tree-level traits. Compared with the spatially continuous map of traits derived at pixel level, the ITC-based trait maps showed additional between-crown gaps. For morphological traits, the linear correlation coefficients (Pearson ρ) of ITC- and 3-m pixel-based H_95_, LAI and FHD were 0.95, 0.65 and 0.62, respectively (Supp Fig. S5). The histograms of traits over the whole study area showed the median values of pixel-based H_95_ and FHD (13.54 m and 0.95) were higher than the ITC-based values (13.25 m and 0.82), while the median value of pixel-based LAI (2.60) was lower than the ITC-based LAI (2.96) (Fig. 5). The histograms also displayed different shapes for ITC- and pixel-based morphological traits. For instance, ITC-based FHD indicated a bimodal distribution, while pixel-based traits showed a smoother (continuous) bell-shaped pattern.

**Fig. 3.**
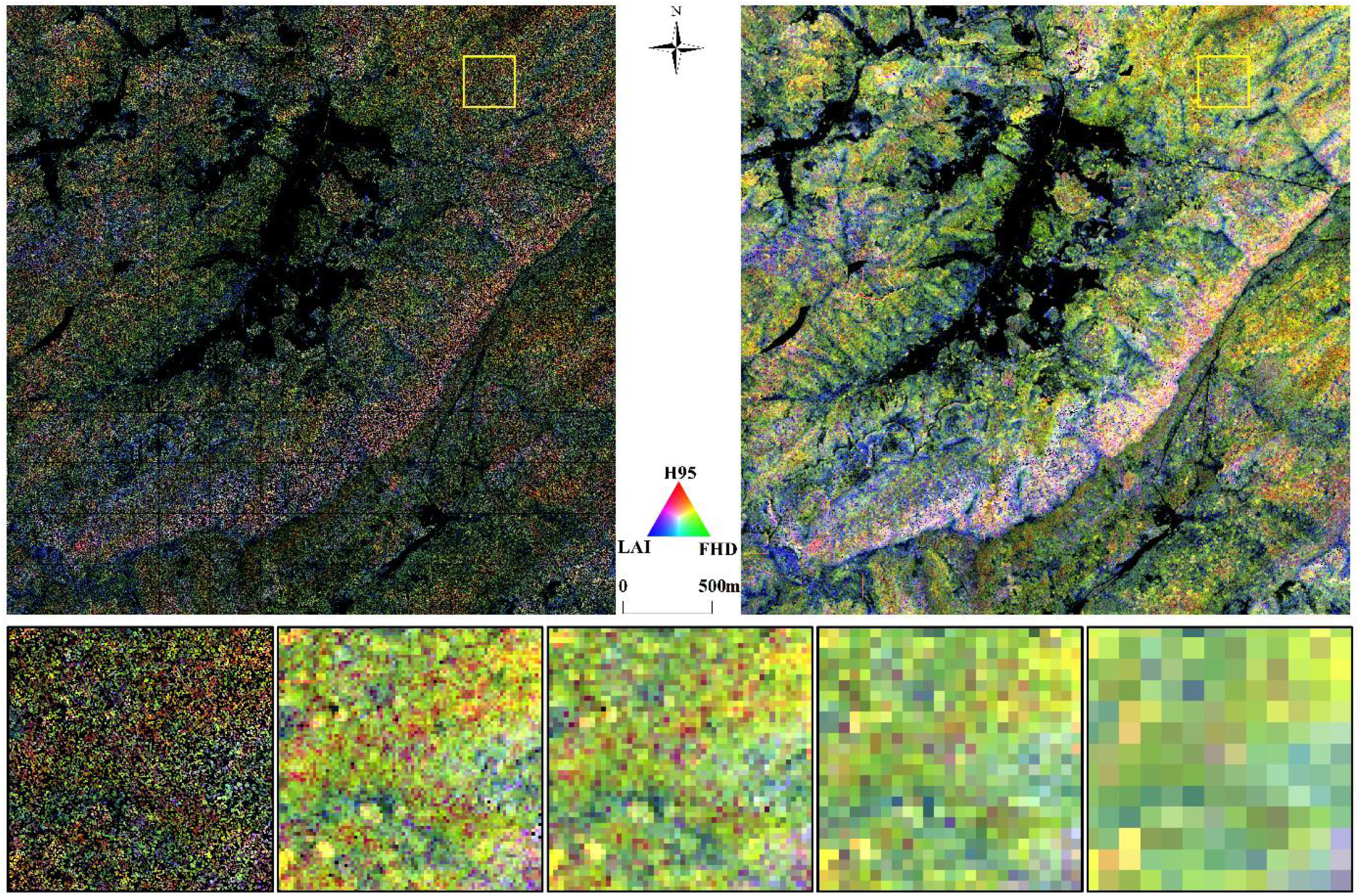
Spatial patterns of morphological traits at individual tree (top left) and 3-m pixel (top right) levels, showing RGB color composites of 95th quantile height (H_95_, red), foliage height diversity (FHD, green) and leaf area index (LAI, blue). The yellow square represents a 250 × 250 m subregion (ITC (bottom far left), and pixel size at 3 m, 5 m, 10 m and 20 m, bottom second left to right).

**Fig. 4.**
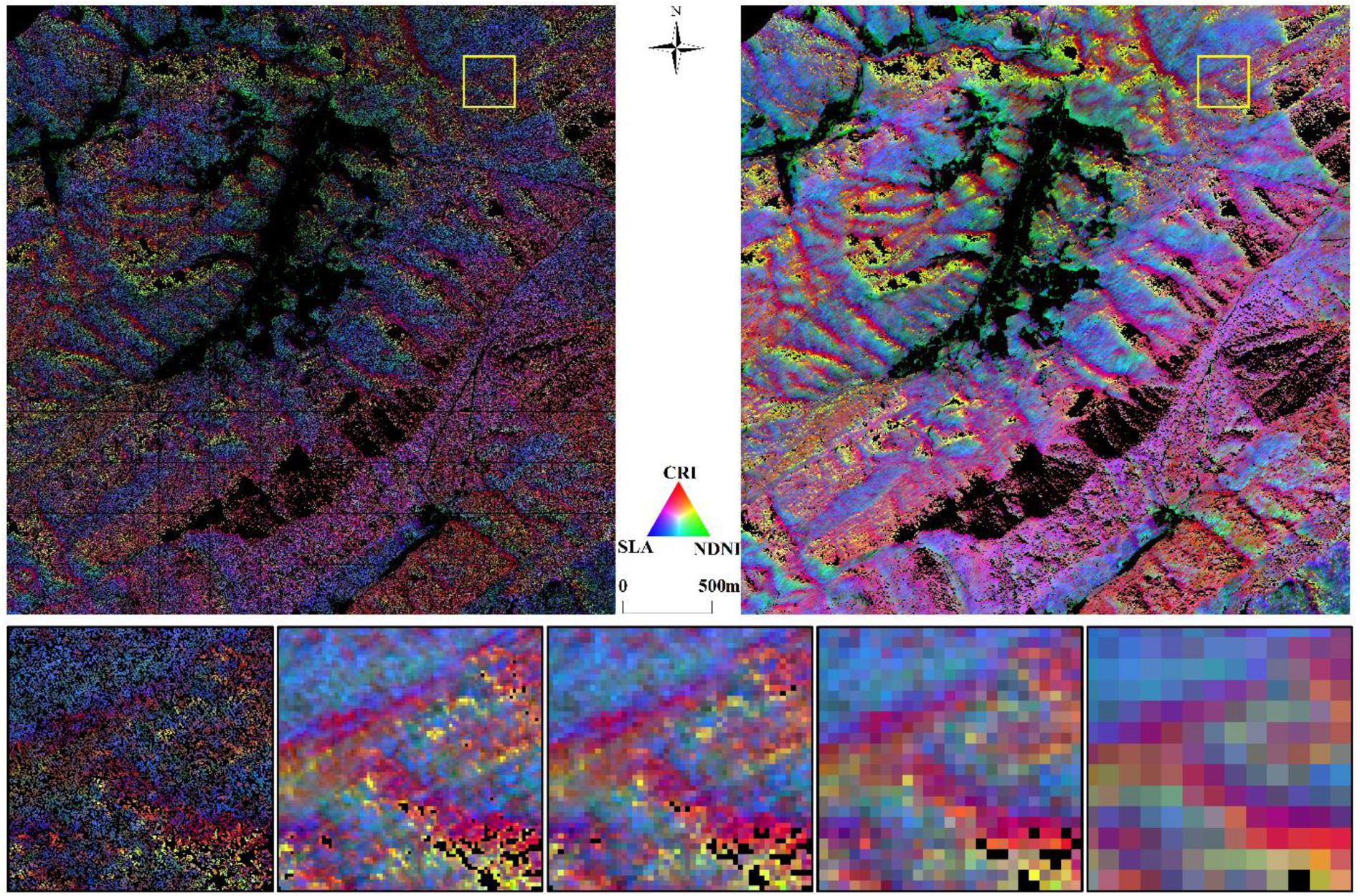
Spatial patterns of physiological traits at individual tree (top left) and 3-m pixel (top right) level, showing RGB color composites of leaf carotenoids (CRI, red), leaf nitrogen (NDNI, green) and specific leaf area (SLA, blue). The yellow square represents a 250 × 250 m subregion (ITC (bottom far left), and pixel size at 3 m, 5 m, 10 m and 20 m, bottom second left to right).

**Fig. 5.**
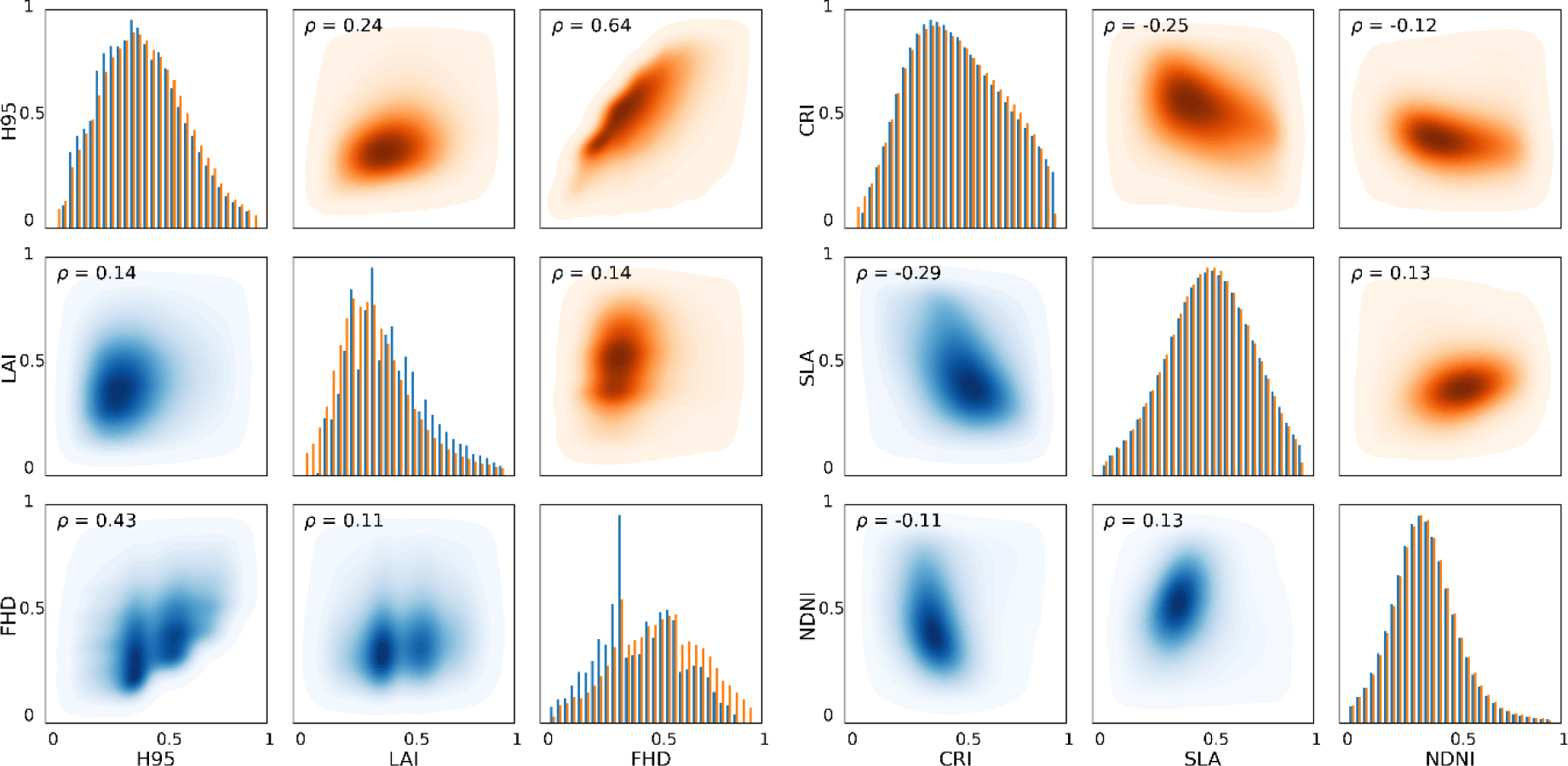
Distribution histograms and correlation matrix of morphological (left) and physiological (right) traits derived from ITCs (blue) and 3-m pixels (orange) over the whole study area (see Supp Fig. S6 for the subregion). Note that patterns of the density scatter plots are mirrored due to the switch of trait axes and the trait values in the scatter plots are linearly scaled from 0 to 1 using a min-max normalization. Darker color indicates higher density of points; Pearson ρ values indicate the trait–trait correlation (p < 0.001) over the whole area.

For physiological traits, the correlation coefficients of ITC- and 3 m pixel-based indicators of carotenoids, SLA and nitrogen were 0.93, 0.93 and 0.91, respectively (Supp Fig. S5). The ITC- and pixel-based physiological traits showed more similar distributions and their histograms indicated similar median values than was the case for the morphological traits (Fig. 5). Analyses of trait–trait relationships showed that trait correlations generally held between ITC- and pixel-based estimates (Fig. 5). For example, H_95_ and FHD were positively correlated for both ITCs (ρ = 0.43) and 3 m pixels (ρ = 0.64), while CRI and SLA were negatively correlated (ρ = −0.29 for ITCs and ρ = −0.25 for 3 m pixels).

### 4.2. Trait distribution and variation at different grain

To illustrate these trait distribution differences for a specific forest community, we chose a 250 × 250 m area of mixed forest and compared ITC- and pixel-based traits at different resolutions (Fig. 6). For morphological traits in this subregion, both K-S test and Mann-Whitney U rank test showed that the ITC- and pixel-based morphological traits were from significantly different distributions (p < 0.001 for all pixel sizes). The pixel-based approach yielded higher H_95_ and FHD and lower LAI than the ITC-based approach (ITC_median_: H_95_ = 16.22, LAI = 2.75, FHD = 0.91), consistent with the results for the whole study area. The IQRs of ITC-based traits (ITC_IQR_: H_95_ = 3.89, LAI = 1.27, FHD = 0.46) were larger than those of pixel-based traits, indicating more variability in the middle half of ITC-based traits. These differences in ITC- and pixel-based morphological traits increased when we increased the pixel size from 3 m to 20 m, and the distributions of pixel-based traits tended to become more clustered. In the subregion, the ITC-based FHD also showed a bimodal distribution, while the histogram of FHD at 3 m pixel size was unimodal and left-skewed. In contrast, for physiological traits, the histogram distributions were quite similar for traits derived from ITCs and 3-m pixels (p > 0.2 in the K-S test and Mann-Whitney U rank test). The differences in physiological traits increased for larger pixels, presumably due to the information mixture when these data were spatially aggregated.

**Fig. 6.**
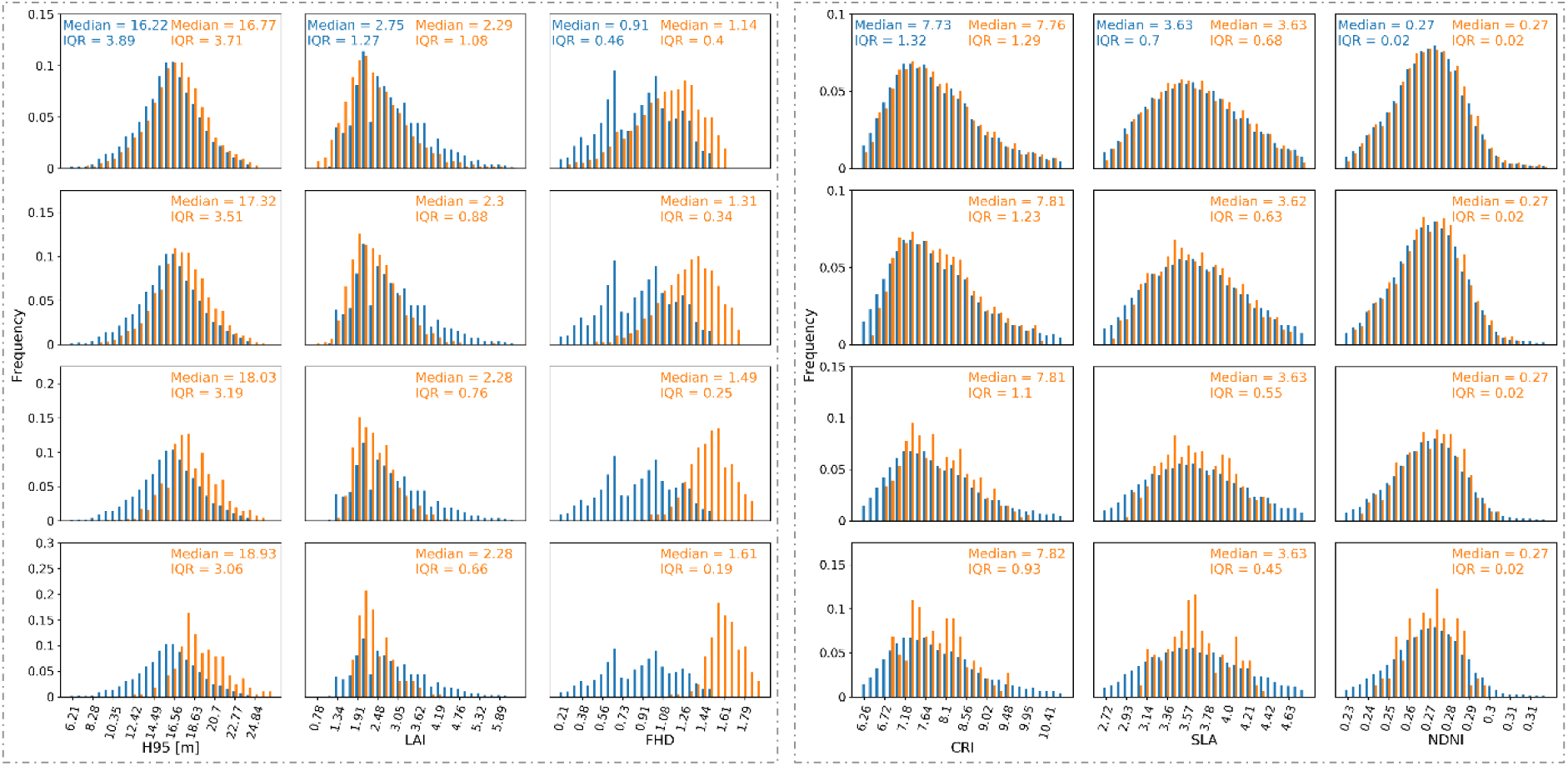
Frequency histograms of morphological (left) and physiological (right) traits derived from ITCs (blue) and pixels (orange) at the 250 × 250 m subregion. The pixel sizes are 3 m, 5 m, 10 m and 20 m from top to bottom panels. Median and IQR values indicate the center and spread of the trait distributions.

The between-unit variation in functional traits at different pixel sizes relative to 3-m pixel-based traits showed that less relative variability in traits could be explained by larger pixels (Fig. 7). For example, the variation in morphological traits (H_95_, LAI and FHD) derived from 10 m pixels could still measure 88.17%, 81.54% and 70.11% of the variance in the whole study area compared with the 3 m pixel size. With the pixel size increased to 25 m, only 76.43%, 75.72% and 56.03% of the morphological trait variability at 3 m pixel level can be captured. It can also be observed that the trait variance decreased a lot when pixel size increased at the small scales (e.g. 3–50 m), and the decrease trends gradually flattened out at larger scales.

**Fig. 7.**
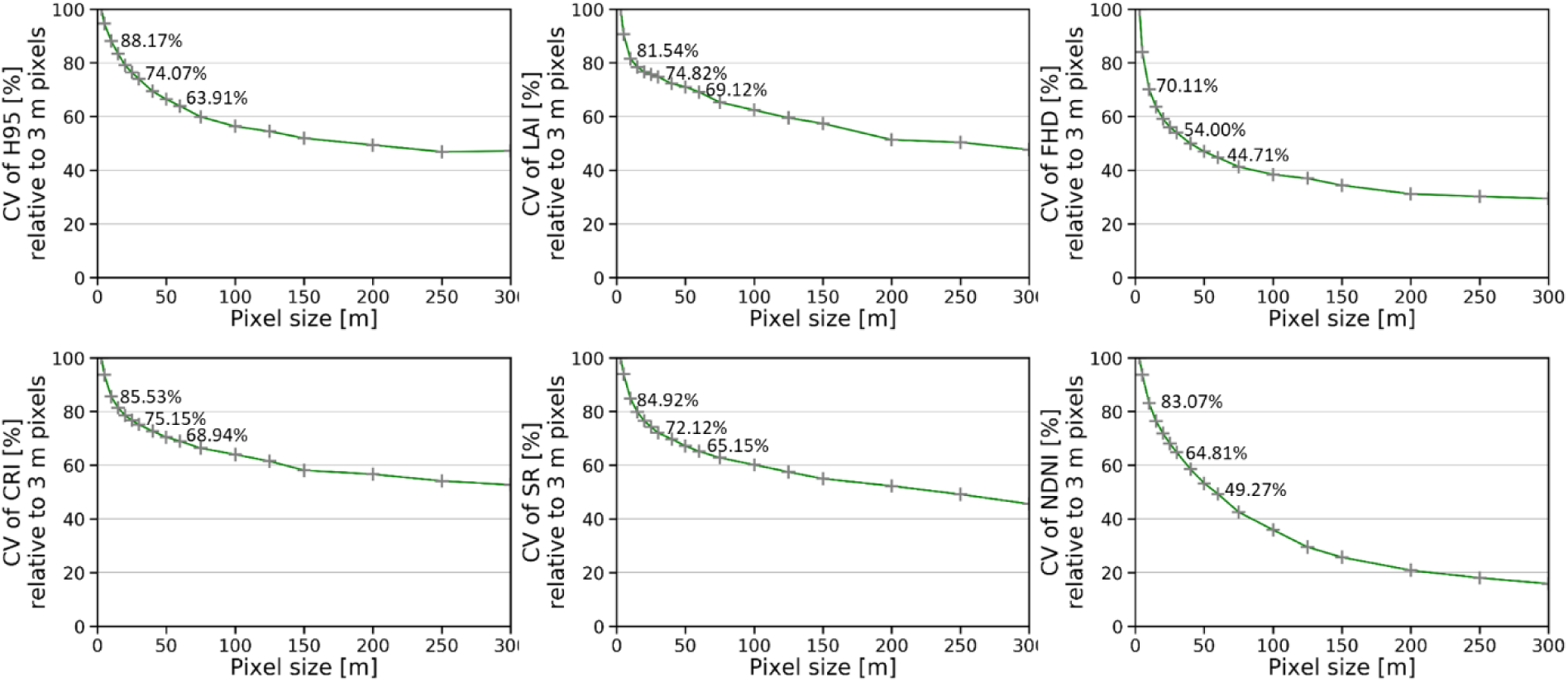
One-dimensional scale analysis for morphological (top) and physiological (bottom) traits over the study area. The various scales include pixel sizes of 3 m, 5 m, 10 m, 15 m, 20 m, 25 m, 30 m, 40 m, 50 m, 60 m, 75 m, 100 m, 125 m, 150 m, 200 m, 250 m, and 300 m, respectively. The numbers marked (from left to right) in the figures indicated the percentage of the trait variability could be detected at 10 m, 30 m and 60 m pixels compared with the 3 m pixel-based trait variation in the study area.

### 4.3. Functional diversity at changing grain and extent

The maps of morphological richness (Morph.FRic) and physiological richness (Phys.FRic) at 30 resolution are shown in Fig. 8 (see Supp Fig. S7 for maps at 100 m resolution), where the ITC- and 3-m pixel-based functional richness estimates displayed similar spatial patterns. A hump-shaped pattern of Morph.FRic along the elevational gradient of 1124–1922 m was well captured by both ITC- and pixel-based approaches, while high Phys.FRic was observed at medium and high elevations (Supp Fig. S8). The pixel-wise correlation coefficients of ITC- and 3-m pixel-based functional richness maps (0.77 and 0.87 for Morph.FRic, 0.86 and 0.92 for Phys.FRic at 30 m and 100 m resolutions, p < 0.001) also indicated good consistency of the two approaches.

**Fig. 8.**
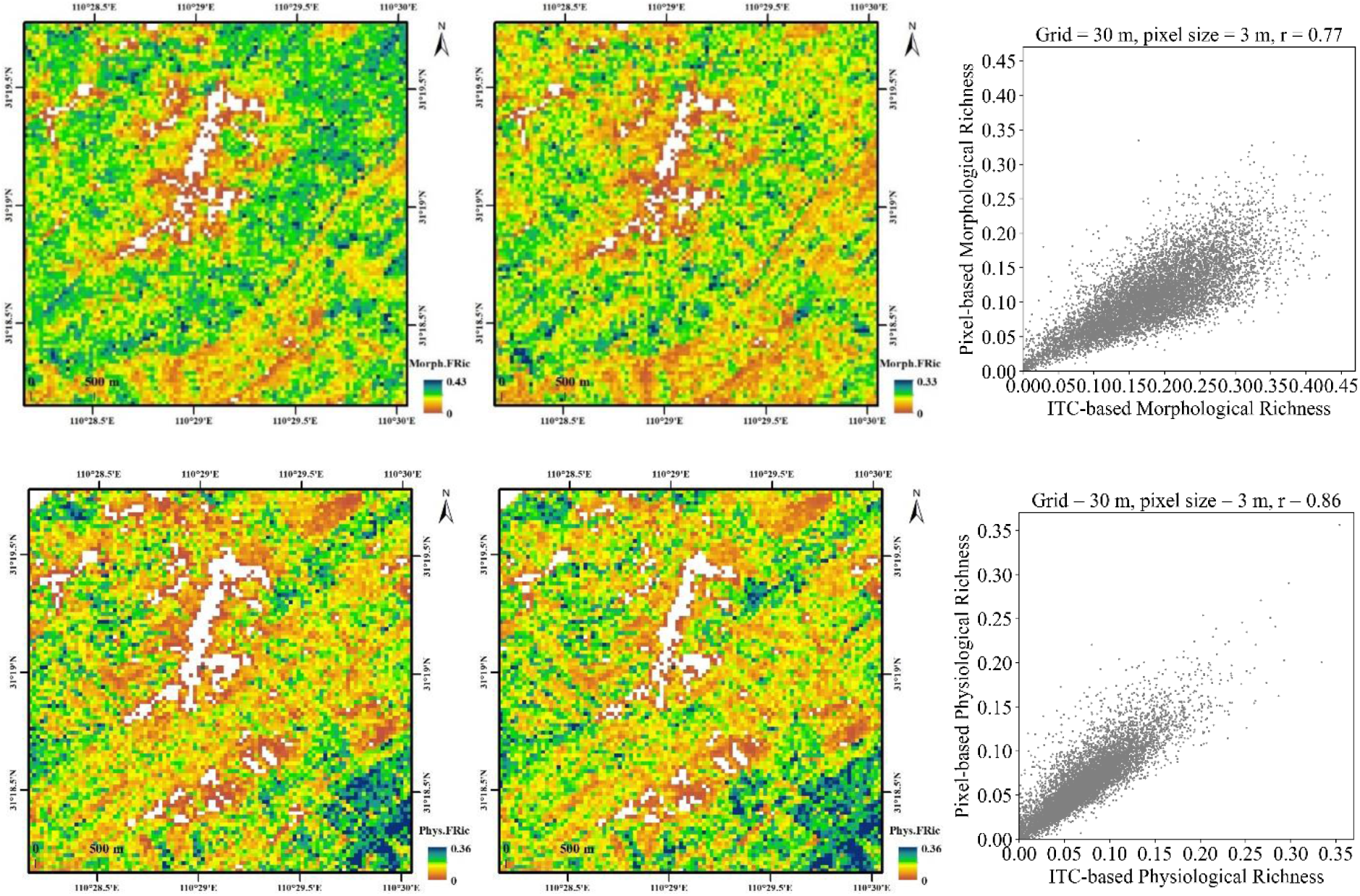
Spatial patterns of morphological (upper) and physiological (lower) functional richness at 30 m resolution based on traits at individual tree (left) and 3 m pixel level (middle) and the corresponding scatter plots of ITC- and pixel-based functional richness measures (right).

The comparisons of ITC- and pixel-based functional richness measures at changing grain and extent showed that for small pixel sizes (e.g. 3 m and 5 m), ITC- and pixel-based functional richness measures correlated very well (Fig. 9). The Pearson correlation coefficients (r) of ITC- and 3 m or 5 m pixel-based functional richness gradually increased and leveled off with increasing neighborhood extents (e.g. r > 0.8 for physiological richness at 24 m radius extent and larger). With increased grain (pixel size), the correlations of ITC- and pixel-based functional richness decreased (see Supp Fig. S9 as an example). Besides, the correlations between ITC- and pixel-based morphological richness were very low and even non-significant for pixel sizes over 15 m. In contrast, ITC- and pixel-based physiological richness still showed relatively good correlations at large grain and extent (e.g. r = 0.65 for ITC- and 15-m pixel-based physiological richness at 150 m radius extent).

**Fig. 9.**
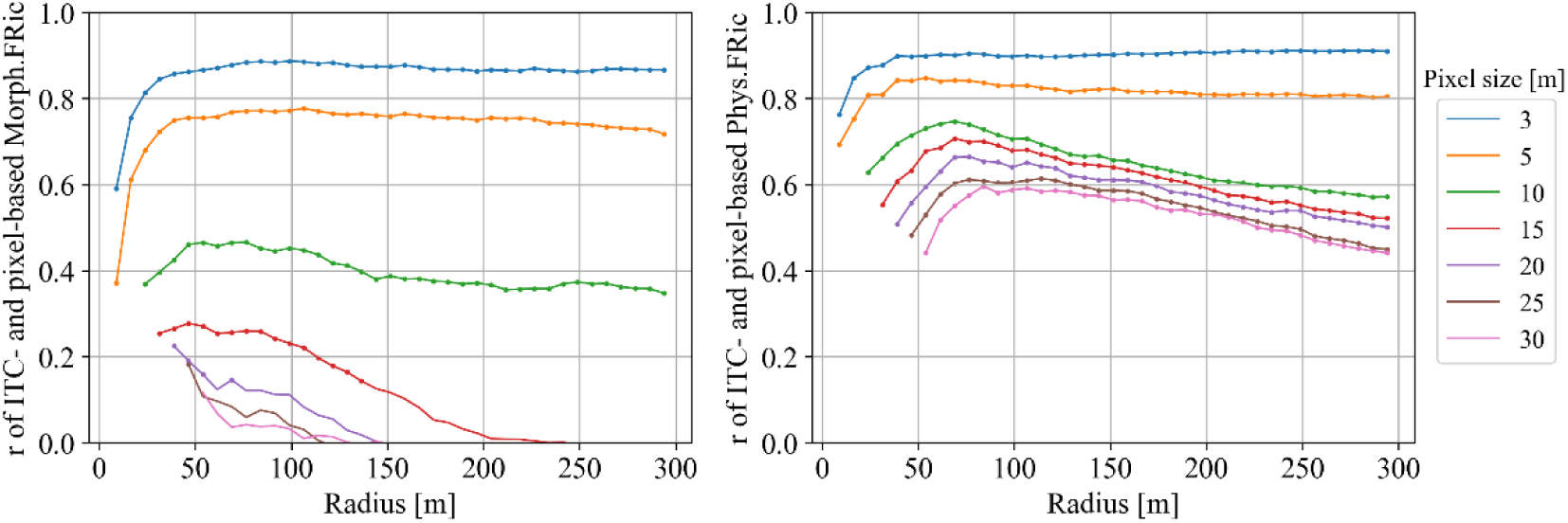
Pearson correlation coefficients (r) between individual tree-based and pixel-based functional richness (FRic) at changing grain and extent for morphological (left) and physiological (right) traits. The colors represent different pixel sizes used in the pixel-based approaches; the dots indicate that the correlations between ITC-based and pixel-based functional richness were significant (p < 0.0001). The radii (9–294 m) indicated different spatial extents of concentric circles surrounding the 1000 sample points for functional richness calculation.

The functional richness–area curves showed that both ITC- and pixel-based functional richness indices increased with area logarithmically (Fig. 10; Supp Fig. S10). For both ITC- and pixel-based approaches, the spatially randomly distributed functional traits (null models) lead to relatively higher functional richness values than remotely-sensed functional richness estimates (see Supp Fig. S10 for mean and standard deviations), which could reveal the prevalence of local trait convergence in our study area. Moreover, ITC-based functional richness increased faster with area at small neighborhood scales than pixel-based estimates, especially for morphological richness, indicating that increased within-community diversity could be better captured by the ITC-based approach.

**Fig. 10.**
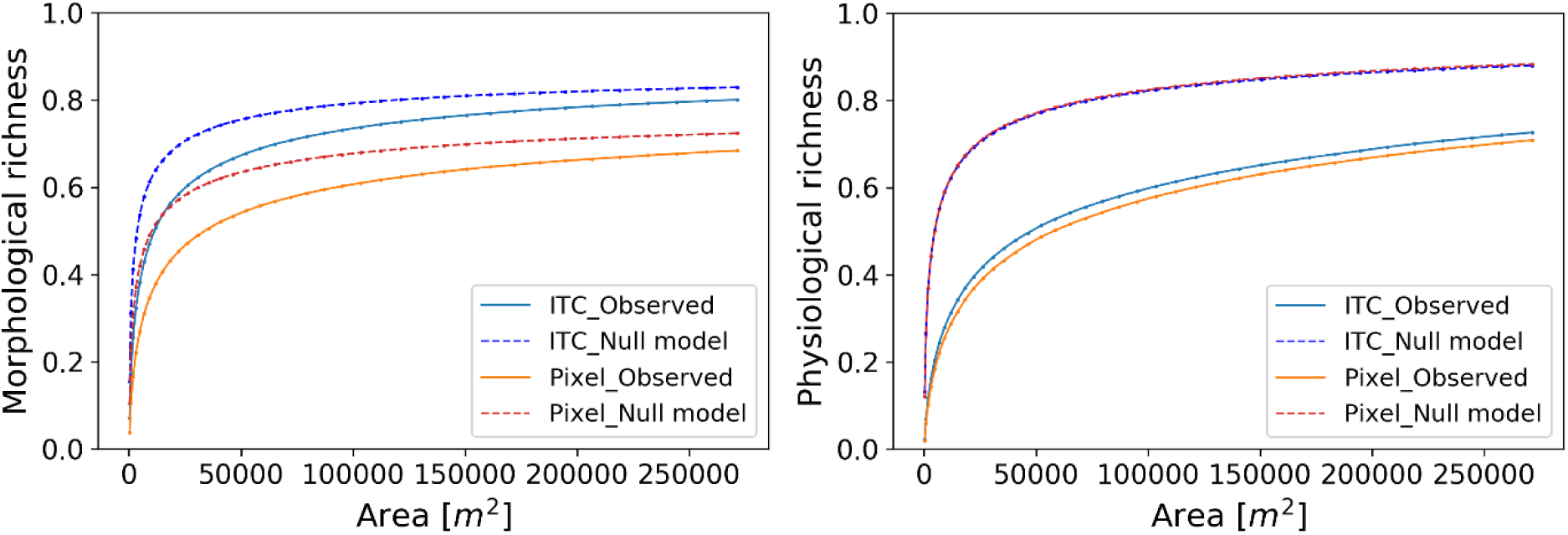
Morphological (left) and physiological (right) functional richness–area relationships based on ITC- and pixel-based traits. Solid lines correspond to the observed relationships from remote sensing; dashed lines depict the relationship between functional richness and area for spatially randomly distributed traits (null models).

## 5. Discussion

### 5.1. Why are the trait distributions different between ITCs and pixels?

Our study shows the potential of converting airborne remote sensing data into ecological information at ITC- and pixel-level and demonstrates the associations and differences among ITC- and pixel-based estimates of functional traits. This comparison is important, because the ITC-based approach assesses individual-, genotype- or species-level functional traits as assembled in databases such as TRY (Kattge et al., 2020), whereas the pixel-based approach generally corresponds to plot-level measurements of community-level functional trait means, which may contain parts or entire individuals of a single or several species. Furthermore, ITCs include only potential within-crown gaps, while pixels also include between-crown gaps. Finally, pixel-based approaches use crown area rather than tree density as a measure of abundance when aggregating to community-weighted means. These reasons may cause the differences in trait distributions obtained with the two approaches.

H_95_ at pixel level applies a 95% quantile filter to height information of all point clouds, which results in higher values when a mixture of taller and lower trees occurs in the pixel. The LAI estimation from the Beer–Lambert law shows smaller values at pixel than at ITC level, which could be explained by the larger gap probability at pixel level (i.e. including multiple trees) than that at individual tree level, and the smaller the fractional crown coverage (*f_cover_*), the larger the difference between ITC- and pixel-based LAI (Supp Text S2). This finding is consistent with previous studies that regarded the pixel-based LAI as the product of the *f_cover_* and average LAI of individual trees in the grid cell (Jiang et al., 2021; Morsdorf et al., 2006; Yang et al., 2019). FHD displays higher values at pixel level, which is also expected to increase with stand height and more even foliage arrangement when multiple trees are covered by a pixel. The choice of spatial unit and ITC- or pixel-based retrieval approach affects the interpretation of the derived morphological traits. The ITC-based FHD shows a bimodal distribution, which might be caused by the different crown architectures of conifer vs. broad-leaved tree species. Pixel-based FHD reflects vertical vegetation strata at the stand or plot level, especially if pixel sizes are larger than the average crown size. For high-resolution pixel, the differences in morphological traits derived from the two approaches are relatively small. Thus, if ITCs are unavailable, an approach using pixels smaller or about the size of ITCs can still capture a large part of trait variation among relevant biological units such as individuals, genotypes and species. However, it should be noted that morphological traits derived from ITC- and pixel-based approaches became more different with increasing pixel size, which indicated different structural features when either individual-tree level or plot level are considered.

For physiological traits, we observed similar spatial patterns and histogram distributions between ITCs and 3-m pixel estimates. The reason might be that biochemical constituents primarily contribute to the spectral response of sunlit pixels within canopy crowns. And both ITCs and synthetic pixels approximating crown size were dominated by sunlit canopy pixels since the shadow and non-forest masks were applied to original images before aggregating. However, as expected, the correspondence between ITC- and pixel-based physiological traits decreased with increasing pixel size (Fig. 6). Unlike the ITC-based approach that only integrates the signal of canopy reflectance inside crown boundaries, larger pixels were more likely to cover multiple crowns and species and include not only tree crowns but also understory and soil in gaps. This then can affect the underlying relationship between foliar traits and their spectral signal (Marconi et al., 2021; Stenberg et al., 2016). Besides, the pixel reflectance is influenced by the overall canopy structure (e.g. spatial arrangement of trees) and multiple scattering driven by vegetation structure as well as mutual shadowing between trees and the fraction of soil (Li and Strahler, 1992, Zeng et al., 2009). This may lead to reflectance differences between data from tree crowns and vegetation canopies (Forsström et al., 2021), thus influencing the distribution of hyperspectral reflectance-based physiological traits. The availability of sub-meter resolution imagery and LiDAR data allows a better characterization of target trees, removing shadow and background contributions (Asner et al., 2007), and capturing the spectral variability within and among crowns. Therefore, by improved ITC segmentation and extracting the sunlit pixels within the crowns, the ITC-based approach can provide robust physiological trait estimates comparable to in-situ observations.

### 5.2. Effects of grain and extent on trait variability and functional diversity

Aggregating fine-scale airborne data allows us to investigate how much trait variation could be captured by coarser pixels, namely, to what level functional diversity can be measured using community-level traits. For both morphological and physiological traits, we noticed that the trait variability explained by pixels decreased with increasing pixel sizes (3 to 300 m, Fig. 7). The trait variability decreased obviously at small grain sizes until around 30 to 50 m, which might indicate that the fine-scale trait variation information of species or individuals within local forest communities was largely lost for pixels at these grains. Similarly, Helfenstein et al. (2022) found a substantial decrease in the variance of physiological traits in a temperate forest when aggregating 6-m pixels to 20-m pixels. However, for scale-aware global forest biodiversity monitoring, the generality and scale thresholds for remotely-sensed trait diversity in different ecosystems need to be further investigated.

Functional richness is sensitive to the range and number of individual points (ITCs or pixels at different resolutions) in the trait space. The interpretations of functional richness should carefully consider the semantics (observation object and meaning of traits) and distribution of the involved traits. With increased grain (pixel size), the correlations of ITC- and pixel-based functional richness decreased. The ecological reason might be that ITC-based functional diversity measures reflect phenotypic and genetic variation among individuals, genotypes and species, which here seemed to be particularly large for morphological traits, whereas pixel-based measures reflect the environmental variation among pixels, which show decreasing overlap with individuals, genotypes and species as pixel size increases. Besides, the correlations between ITC- and pixel-based functional richness measures were relatively lower at small neighborhood extents when only a few pixels were involved in the calculation. With increasing observation extents, we found an interesting pattern for physiological richness measures based on pixels larger than 10 m: the correlation of ITC- and pixel-based functional richness first increased but then decreased slightly. A potential reason for the decreased correlation at larger extents might be more influence of soil or other non-canopy information for pixel-based traits due to less strict filtering conditions than ITCs. For example, more pixels near the forest edge might be included in the aggregation of larger pixels, while small pixels or ITCs would be less influenced because ITCs and synthetic pixels that had more than half non-forested or shaded areas were excluded.

The multi-scale comparison revealed how much of the ITC-level trait diversity can be observed using different resolution pixel data, and indicated the potential of using medium-high resolution spaceborne data (such as Sentinel-2, PRISMA) to assess forest functional diversity over large areas. These satellite imagery generally has a pixel resolution larger than most tree canopies and the derived functional traits will thus correspond to the community-level traits. The resolution of airborne data is usually finer than the crown size, thus could be used to extract individual-level traits. The 30 m or 100 m resolution functional richness derived from traits of ITCs or high-resolution pixels can represent the α-functional diversity. When upscaling functional traits to larger pixels (e.g. Sentinel-2), the functional richness likely captures both α- and β-diversity, indicating the local turnover of functional traits. Generally, the remote sensing detection of α-diversity depends on plant size and image resolution. For larger pixels that capture differences in plant community composition (e.g. 30 m), β-diversity measures quantitating differences between vegetation inventory plots /pixels can be a promising tool for large-scale remote sensing of biodiversity (Leutner et al., 2012; Schweiger and Laliberté, 2022).

### 5.3. Advantages and limitations of ITC- and pixel-based approaches

Our findings demonstrate how forest functional diversity could be assessed from remotely-sensed functional traits with either ITC- or pixel-based approaches. Recognizing the advantages and the limitations of these two approaches to trait retrieval and data interpretation is important for choosing a suitable approach in different contexts. Many remote sensing-based trait estimations are computed at the pixel level, from medium- to high-resolution airborne or spaceborne data (Coops et al., 2016; Hauser et al., 2021; Schneider et al., 2017), where pixels are regarded as composites of individuals, populations or communities. Although pixel values rarely represent species or individuals, the pixel-based approach has the advantage of mapping community-weighted traits continuously and monitoring functional diversity over large areas and at a global scale with abundantly available active and passive remote sensing data (Jetz et al., 2016; Wang and Gamon, 2019; White et al., 2016). Pixel-based approaches combined with multi-temporal satellite data can monitor vegetation dynamics and potentially reveal phenological characteristics of plant traits efficiently, which can also provide additional variables for inferring biodiversity (Rocchini et al., 2016; Rossi et al., 2021).

Trait mapping and functional diversity monitoring for millions of individual trees may be achieved by combining airborne LiDAR and imaging spectroscopy data within crown boundaries (Zheng et al., 2021). The main advantage of ITC-based trait estimations is that they better allow the analysis of trait variation related to the biological units that cause this variation due to their different genetic and phenotypic characteristics and responses to local environmental conditions. The ITC-based approach allows us to directly account for variation between individuals and thus makes it possible to separate overall trait variation into contributions from species and higher taxonomic or phylogenetic groups and environmental variates over spatially contiguous areas (Guillén-Escribà et al., 2021). ITC-based approaches are therefore preferred in ground-based surveys, against which airborne surveys using ITCs can then be compared (Dalponte and Coomes, 2016). Furthermore, ITC-based trait estimates minimize “contamination” from non-biological objects (e.g. bare soil, stones, etc.) or biological objects (e.g. ground vegetation in gaps between tree crowns) that are not to be included in a particular biodiversity assessment.

For morphological traits, ITC-based approaches can capture the detailed architectural information of each detected tree and provide insights into the structural variation such as size– density relationships and forest dynamics at the organismal level that cannot be achieved for pixels (Baruffol et al., 2013; Duncanson and Dubayah et al., 2018). For physiological traits, previous studies reported that aggregating information to the ITC level could improve the accuracy of trait estimations (Marconi et al., 2021) and tree species discrimination (Torabzadeh et al., 2019), because individuals can not represent multiple species while pixels can. It has also been reported that physiological indices at ITC level perform better in monitoring water stress and fruit quality than coarse pixels affected by soil reflectance and between-tree shadows (Stagakis et al., 2012). Moreover, open-access remotely-sensed ITC datasets and deep learning (Brandt et al., 2020; Weinstein et al., 2021; Yang et al., 2022) could support individual-level ecological studies at large scales, opening up additional directions for analyses, such as tree density–diversity relationships and how these affect ecosystem functioning in real-world forests.

Despite advances in the application of airborne remote sensing to retrieve functional traits and resolve ecosystem questions, there remain technical uncertainties and issues which need to be addressed. One limitation of both ITC- and pixel-based approaches is the underrepresentation of sub-canopy information due to the top-down perspective, which means it may be unsuitable for ecological analyses that require accurate characterization of sub-canopy vegetation. For forest diversity research, previous studies from boreal forests to the tropics suggested that remotely sensed diversity primarily based on the upper canopy layer matched well with diversity estimates from forest inventories (Bohlman, 2015; Ma et al., 2019). Another issue requiring careful consideration is the uncertainty in ITC detection. In our subtropical forest site, LiDAR data allowed us to match 83.4% of trees with higher confidence in detecting taller trees (Supp Fig. S1). However, CMH-based segmentation algorithms tend to under-segment overlapping canopies and lower-canopy strata (Liu et al., 2017; Zheng et al., 2021). Besides, misalignment between field and algorithmically delineated crown polygons was reported in a recent study, but it showed that the cascading impacts on trait estimation were generally small (Marconi et al., 2021).

Therefore, it is important to consider how this uncertainty propagation will influence the interpretation of data in light of specific research questions, and whether the advantage of the individual perspective outweighs errors in crown delineation. Fortunately, a number of questions and applications in ecology are primarily influenced by large canopy trees, for instance, overstory trees play much more important roles in carbon and nitrogen cycling than understory trees (Moore et al., 2007). In summary, whether ITC- or pixel-based approaches are more suitable or relevant depends on the specific research question. For example, the remote sensing assessments of ecological processes and functions involving individual tree interactions and plant demography (e.g. tree growth and mortality, complementarity and selection) might benefit from the individual perspective (Worthy and Swenson, 2019; Yang et al., 2018), while for some biogeochemical and biophysical functions (e.g. related to short-term CO_2_ dynamics) at large scales, pixel-based approaches might be sufficient.

## 6. Conclusions

In this study, we compared ITC- and pixel-based approaches for monitoring functional traits and investigated how grain and extent affect trait-based functional diversity using airborne LiDAR and imaging spectroscopy data. Typically, if pixel size became larger than average crown size, the inclusion and averaging of multiple species and between-crown gaps made the difference between the approaches larger, reflecting species- vs. community-level functional trait assessments. With increasing pixel size, less variance in traits can be detected by the pixel-based approach, indicating information loss of the small-scale variability. The consistency comparison of ITC- and pixel-based functional richness measures at changing grain and extent could support identifying the optimal scale (both pixel size and monitoring extent) for effective functional diversity assessment when only coarser pixel data are available. The ITC-based approach is directly analogous to field-based approaches for trait sampling of individuals. Wall-to-wall ITC-level functional traits may provide a new view on individual tree-based ecology at large scales (e.g. ecological processes involving individuals), but very-high-resolution data is a prerequisite. The pixel-based approach can scale up functional traits and diversity to larger areas in a continuous and repeatable way using satellite imagery, although much fine-scale variability will be lost. Whether an ITC- or pixel-based approach is more suitable depends on the spatio- temporal resolution of the data and the relevance of the individual perspective in a specific research question.

Our comparison of the ITC- vs. pixel-based approach could help us understand the linkages and differences between species-level and community-level functional trait assessments. Different ways of data aggregation to ITCs and pixels could also provide insights to upscale functional traits from local to regional and global scales. Currently, airborne data are generally available at regional scales as a complement to ground observations and are essential for capturing fine-scale trait variation in a region. They can be further combined with satellite data to effectively estimate functional traits over large extents and scale-up assessments of functional diversity, by providing more model training data and aggregating to the grain size matching with satellite imagery. Spatial aggregation of high-resolution airborne pixels could increase the representativeness for model training and validation, which is important for accurately predicting plant functional traits and diversity across scales. Therefore, it would be helpful to leverage intermediate-scale airborne data to bridge the scale gap between local ground inventory plots and global spaceborne observations.

## Acknowledgements

This study was supported by the National Natural Science Foundation of China (No. 42071344). The contributions of ZZ, FM, MCS, BS and MES were supported by the University of Zurich Research Priority Program on Global Change and Biodiversity (URPP GCB). The authors thank Zongqiang Xie, Wenting Xu, and Changming Zhao from the Shengnongjia Biodiversity Research Station of the Chinese Academy of Science for their support with fieldwork. We also thank Dan Zhao, Yujin Zhao, Xinfeng Zhao, Liang Zhu, Wenxue Dong, Zilong Yang, Zhugeng Duan, Wenwen Gao and Hong Chi for their assistance with the field sample collection. The acquisition and preprocessing of airborne LiDAR and imaging spectroscopy data were supported by the Shanghai Hangyao Information Technology Cooperation. ZZ acknowledges financial support from the China Scholarship Council (No. 201704910941).

## Appendix A. Supplementary data

Supplementary data associated with this article can be found in the separate supplementary materials.

