## Supplementary Material for "Individual tree-based vs pixel-based approaches to mapping forest functional traits and diversity by remote sensing"

### Contents of this file

Text S1 Individual tree crown (ITC) segmentation approach

Text S2 Difference between ITC- and pixel-based LAI

Figures: Supp Fig. S1 to S10

### Introduction

This supporting information provides two supplementary texts and ten supplementary figures for further illustrating our method and results.

### Text S1 Individual tree crown (ITC) segmentation approach

The ITC segmentation approach, i.e. a morphological crown control-based watershed algorithm, was described in detail in Zhao et al. (2014) and was used and validated in our previous study (Zheng et al., 2021). The processing mainly included the following steps: (i) a morphological closing operator combined with a median filter was applied to the original CHM to fill the invalid value in potential crown areas (Zhao et al., 2013) and limit the treetop detection; (ii) a local maxima algorithm to identify potential individual treetop positions; (iii) these treetops were used as markers to constrain the watershed transform algorithm from over-segmenting and two watershed segmentation routines were combined to delineate the individual tree basins; (iv) a morphological opening operator was applied to optimize the basins to approximate shape of crowns; and (v) a quickhull method was applied to LiDAR points within each watershed segment, and the resulting polygons became the final ITCs.

### Text S2 Difference between ITC- and pixel-based LAI

Based on the relationship between the gap probability at ITC level and pixel level (i.e. including multiple trees):

$$P_{pixel} = f_{cover} \cdot P_{tree} + (1 - f_{cover})$$

where $f_{cover}$ means the fractional crown coverage which is between 0 and 1, 0 denotes there is no tree in the pixel and 1 denotes there are no between-crown gaps in the pixel; $P_{tree}$ means the average gap probability of a single tree.

The difference between pixel and tree LAI can be deduced as follows：

$$P_{pixel} - P_{tree} = (1 - f_{cover})\,(1 - P_{tree}) \quad (0 \leq f_{cover} \leq 1, 0 < P_{tree} < 1)$$

$$P_{pixel} - P_{tree} \geq 0;\;\; P_{pixel} \geq P_{tree}$$

$$LAI_{pixel} \leq LAI_{tree}$$

Pixel LAI ($LAI_{pixel}$) is usually smaller than the LAI at tree level ($LAI_{tree}$) if we directly apply the Beer-Lambert law on LAI estimation using the gap probability, and the smaller the $f_{cover}$ (i.e. the more sparse the trees in the forest are), the larger the difference between the $LAI_{pixel}$ and $LAI_{tree}$.

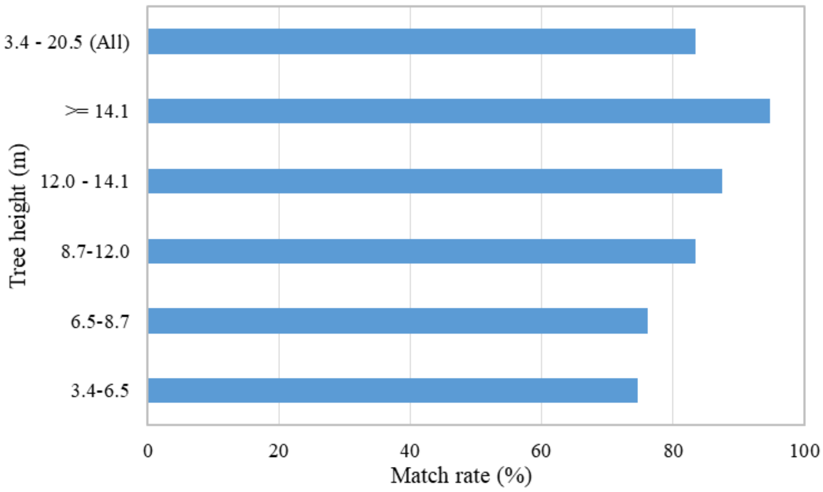


Supp Fig. S1. ITC detection rate for different tree height groups.

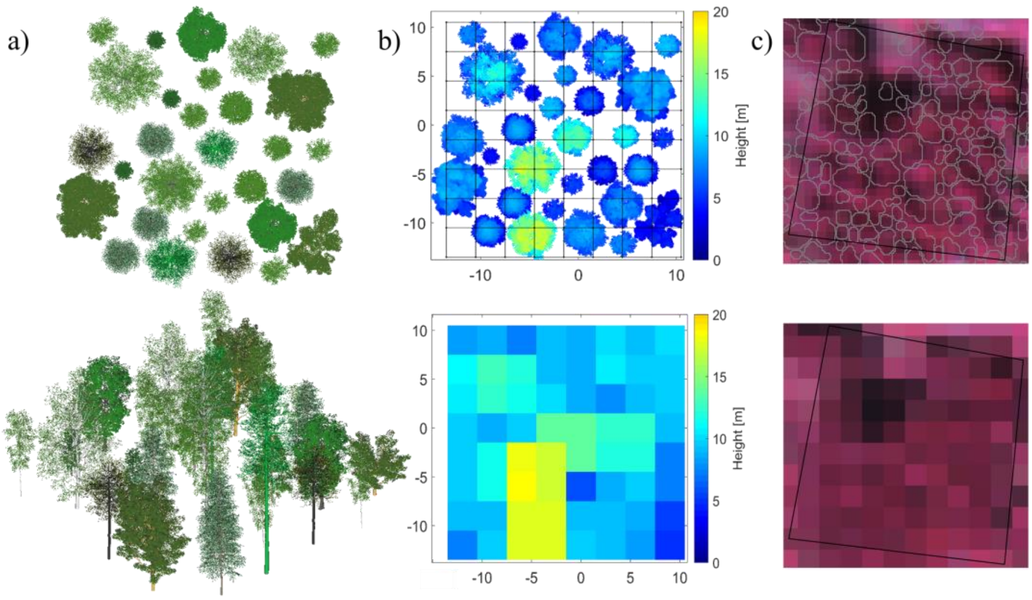


Supp Fig. S2. Schematic diagram of ITC- and pixel-based approaches. (a) a virtual forest scenario (top and front view) created by tree models from OnyxTREE; (b) detailed height information of these tree canopies and aggregated height for 3-m pixels; (c) spatial intersection of the LiDAR-detected crown polygons with hyperspectral imagery to obtain integrated signals for ITCs and directly resampled to obtain signals for 3 m pixels.

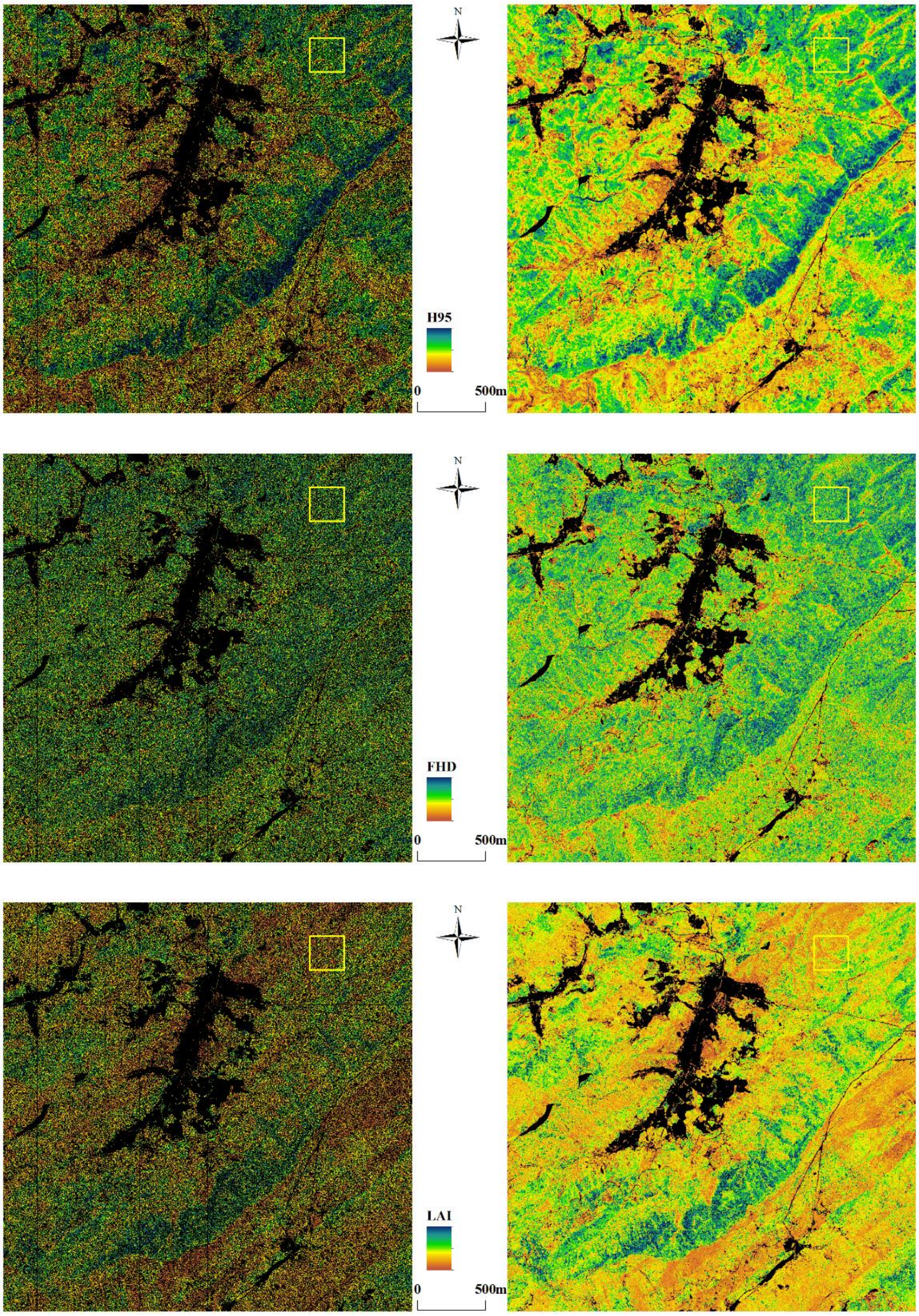


Supp Fig. S3. Spatial patterns of morphological traits at individual tree (left) and 3 m pixel (right) level, the traits are 95th quantile height ($H_{95}$), foliage height diversity (FHD) and leaf area index (LAI) from top to bottom panel.

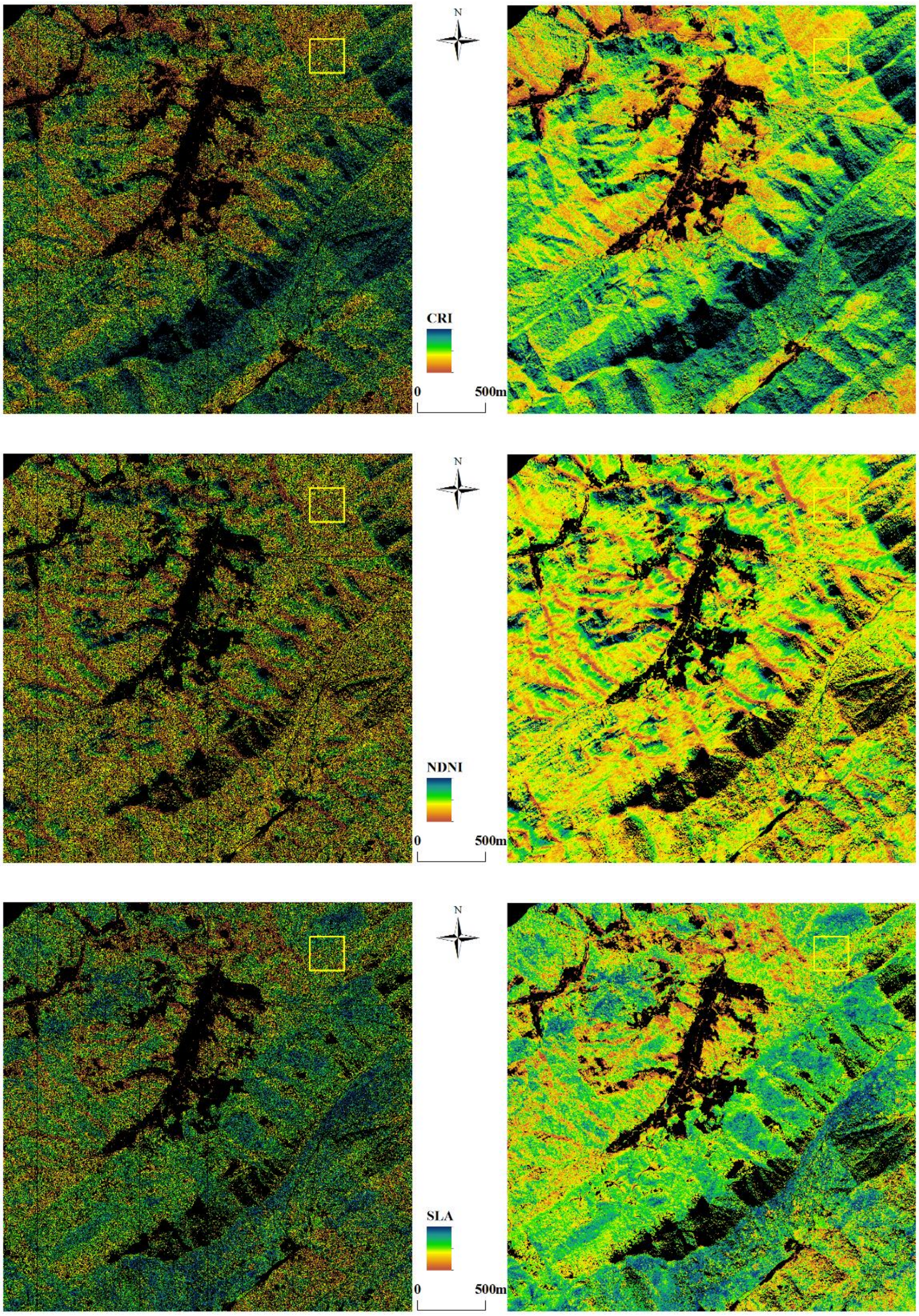


Supp Fig. S4. Spatial patterns of physiological traits at individual tree (left) and 3 m pixel (right) level, the traits are carotenoids (CRI), nitrogen (NDNI) and specific leaf area (SLA) from top to bottom panel.

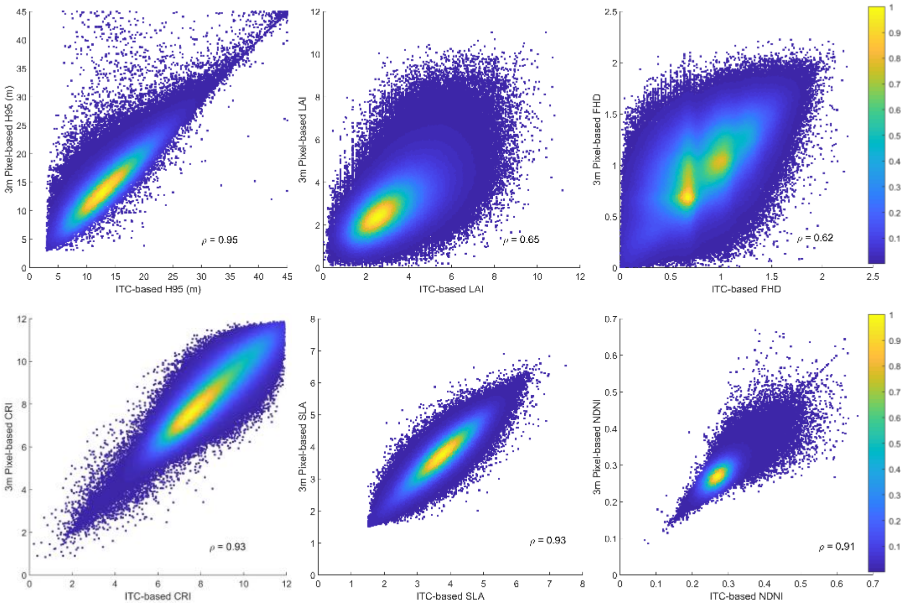


Supp Fig. S5. Pearson correlation coefficients (ρ) between functional traits derived by ITCs and 3 m pixels at corresponding locations. The upper panels are the scatter plots for ITC- and pixel-based morphological traits: 95th quantile height ($H_{95}$), leaf area index (LAI) and foliage height diversity (FHD). The lower panels are the scatter plots for ITC- and pixel-based physiological traits: leaf carotenoids (CRI), leaf nitrogen (NDNI) and specific leaf area (SLA). The colors in the legend indicate density of the scatter data.

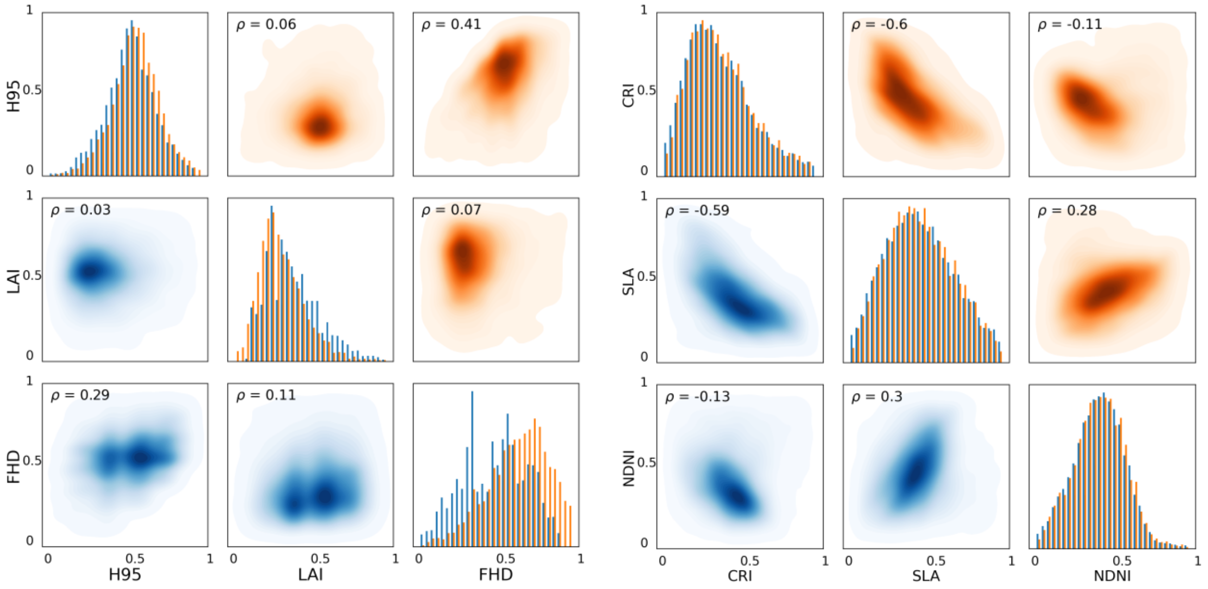


Supp Fig. S6. Distribution histograms and correlation matrix of morphological (left) and physiological (right) traits derived from ITCs (blue) and 3 m pixels (orange) in the 250 × 250

m subregion. Note that patterns of the density scatter plots are mirrored due to the switch of trait axes and the trait values in the scatter plots are linearly scaled from 0 to 1 using a min-max normalization. Darker color indicates higher density of points; Pearson ρ values indicate the trait–trait correlation ($p < 0.001$) over this 250 × 250 m subregion.

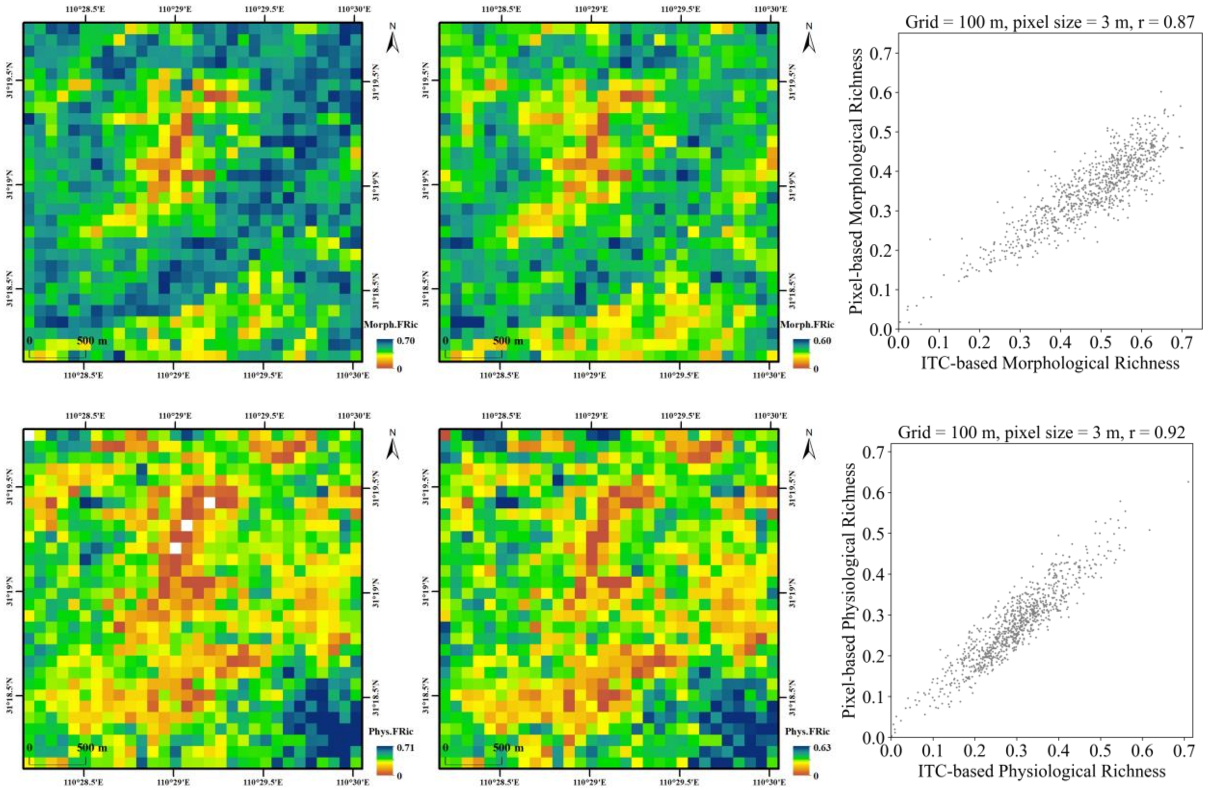


Supp Fig. S7. Spatial patterns of morphological (upper) and physiological (lower) functional richness at 100 m resolution based on traits at individual tree (left) and 3 m pixel level (middle) and the corresponding scatter plots of ITC- and pixel-based functional richness measures (right).

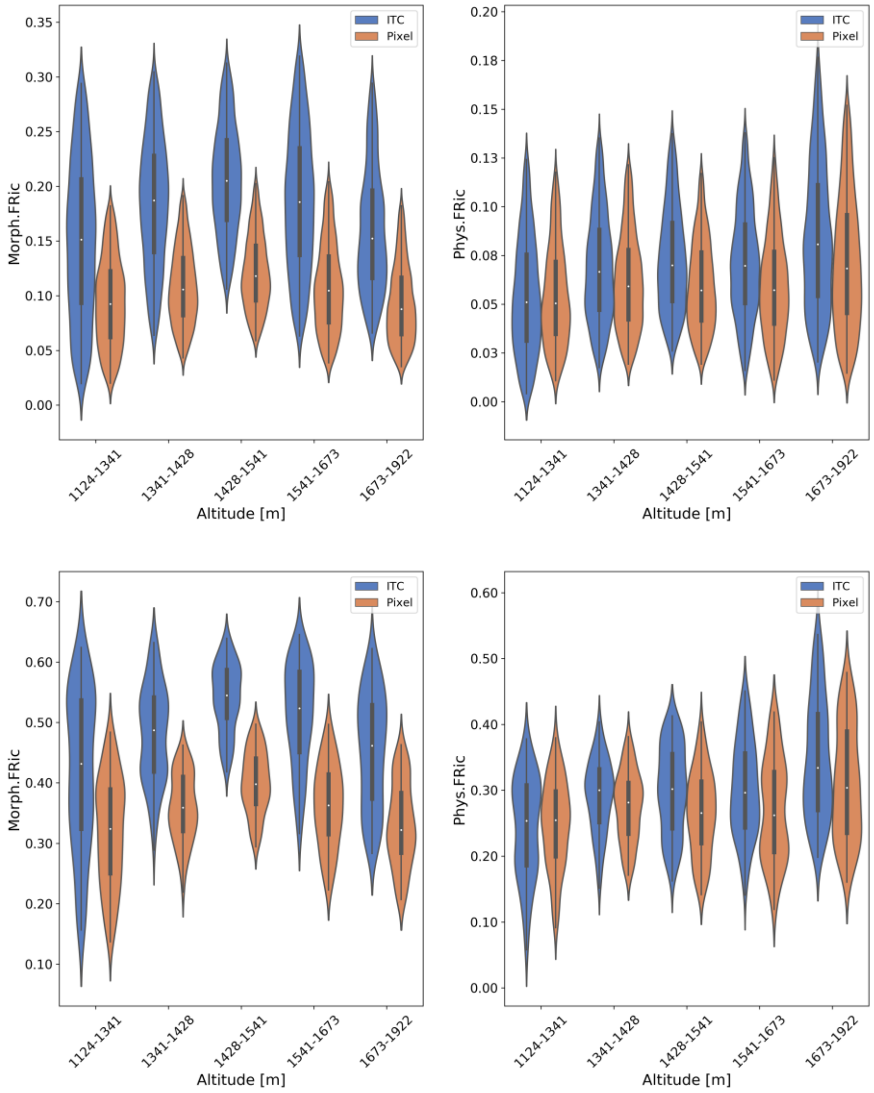


Supp Fig. S8. Violin plots of functional richness indices (between the 5th and 95th percentile) along altitudinal belts. Morphological richness (Morph.FRic, left) and physiological richness (Phys.FRic, right) are shown for a spatial resolution of 30 m (upper panel) and 100 m (lower panel). The median values of functional richness indices in each altitudinal belt are indicated as white points in the miniature boxplots inside the violins. The width of the violin plots represents frequency. The altitudinal belts are based on 20%, 40%, 60% and 80% quantile of the DEM ranging from 1124 to 1922 m in the study area.

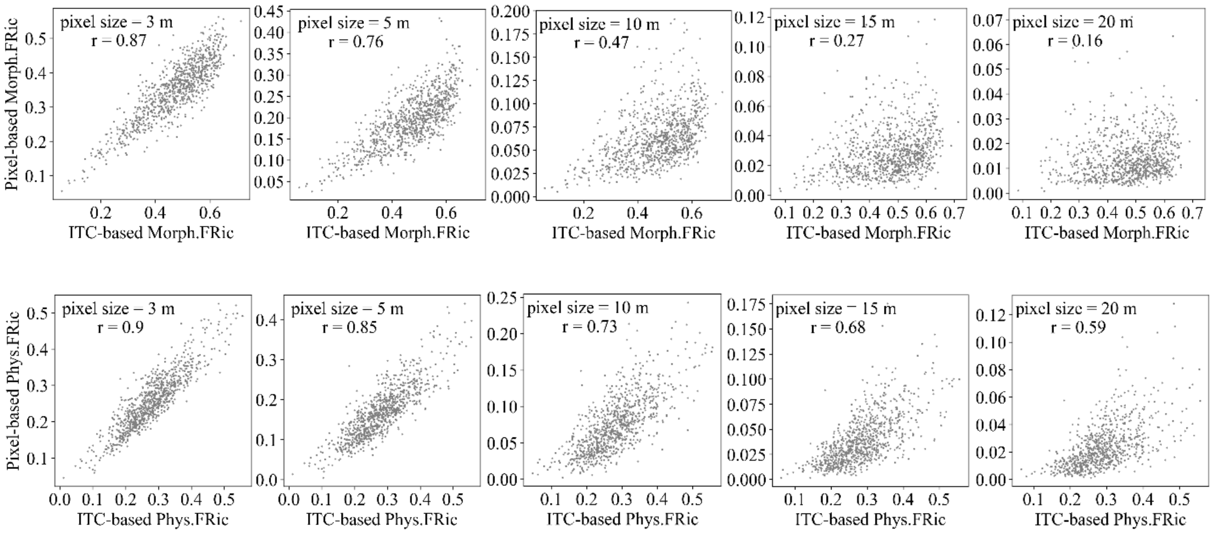


Supp Fig. S9. Scatter plots and Pearson correlation coefficients between functional richness derived by ITCs and pixels at 3 m, 5 m, 10 m, 15 m and 20 m resolution in 54 m radius extent surrounding the sample points. The upper panels are the scatter plots for ITC- and pixel-based morphological richness (Morph.FRic). The lower panels are the scatter plots for ITC- and pixel-based physiological richness (Phys.FRic).

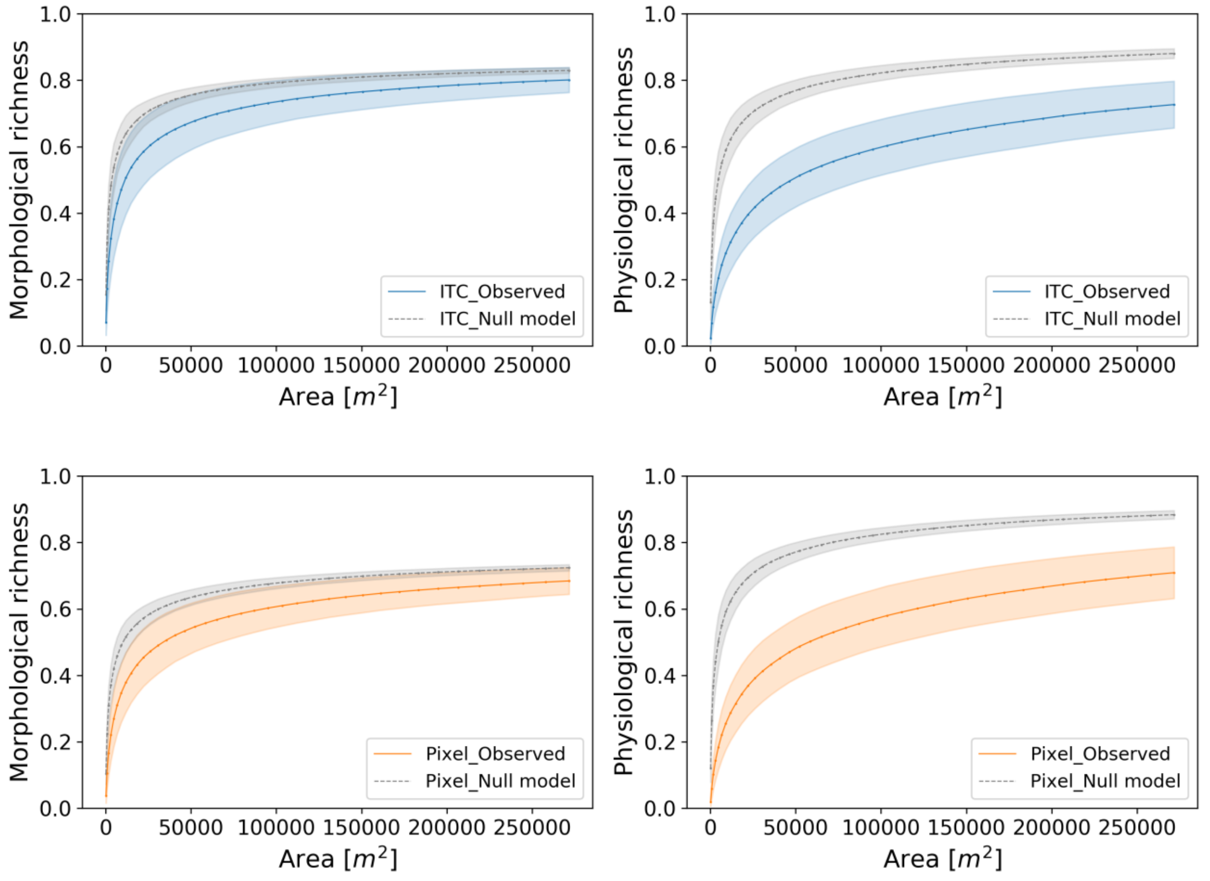

Supp Fig. S10. Morphological (left) and physiological (right) functional richness–area relationships based on ITC- (top) and pixel-based (bottom) traits. Colored solid lines and colored areas correspond to mean and standard deviation of functional richness estimates of all sample central points at each neighborhood extent. Grey lines and shades indicate the mean and standard deviation of functional richness at the null model scenarios of randomly distributed ITCs or pixels.
